# Establishment of a Prognostic Prediction and Drug Selection Model for Patients with Clear Cell Renal Cell Carcinoma by Multi-Omics Data Analysis

**DOI:** 10.1101/2021.08.06.455377

**Authors:** Aimin Jiang, Yewei Bao, Anbang Wang, Xinxin Gan, Jie Wang, Yi Bao, Zhenjie Wu, Bing Liu, Juan Lu, Linhui Wang

## Abstract

**Rationale:** Patients with clear cell renal cell cancer (ccRCC) may have completely different treatment choices and prognoses due to the wide range of heterogeneity of the disease. However, there is a lack of effective models for risk stratification, treatment decision making and prognostic prediction of renal cancer patients. The aim of the present study was to establish a model to stratify ccRCC patients in terms of prognostic prediction and drug selection based on multi-omics data analysis.

**Methods:** This study was based on the multi-omics data (including mRNA, lncRNA, miRNA, methylation and WES) of 258 ccRCC patients from TCGA database. Firstly, we screened the feature values that had impact on the prognosis and obtained two subtypes. Then, we used 10 algorithms to achieve multi-omics clustering, and conducted pseudo-timing analysis to further validate the robustness of our clustering method, based on which the two subtypes of ccRCC patients were further subtyped. Meanwhile, the immune infiltration was compared between the two subtypes, and drug sensitivity and potential drugs were analyzed. Furthermore, to analyze the heterogeneity of patients at the multi-omics level, biological functions between two subtypes were compared. Finally, Boruta and PCA methods were used for dimensionality reduction and cluster analysis to construct a renal cancer risk model based on mRNA expression.

**Results:** A prognosis predicting model of ccRCC was established by dividing patients into high- and low-risk groups. It was found that overall survival (OS) and progression-free interval (PFI) were significantly different between the two groups (p<0.01). The area under the OS time dependent ROC curve for 1, 3, 5 and 10 years in the training set was 0.75, 0.72, 0.71 and 0.68 respectively.

**Conclusion:** The model could precisely predict the prognosis of ccRCC patients and may have implications for drug selection for ccRCC patients.

## Introduction

Renal cancer is a malignant tumor derived from the proximal tubule epithelium in the renal parenchyma(1). It is one of the most common tumors of the urinary system and one of the ten most common tumors in the world(2). The incidence and mortality of renal cancer are increasing yearly. The latest world cancer statistics show that the number of new cases of renal cancer is about 431,288 (2.2%) and the number of deaths is about 179,368 (1.8%) worldwide each year. Clear cell renal cell carcinoma (ccRCC) or Kidney renal clear cell carcinoma (KIRC) is the most common pathological type of renal cancer, accounting for 70-85% of total RCC (3). It is noteworthy that most renal cancer cases are insidious and lack typical clinical symptoms. Therefore, about 20-30% renal cancer patients have metastasized at the initial diagnosis, and 30% patients may have recurrence or metastasis. Renal cancer is not sensitive to traditional radiotherapy and chemotherapy. The current main treatments include surgery, chemotherapy and immunotherapy, but about 30% patients will develop primary or secondary drug resistance. The degree of CD8+ T cell infiltration is positively correlated with a better prognosis in most solid tumors, whereas it is often associated with a worse prognosis in ccRCC (4). It is therefore urgent to gain a better understanding about the heterogeneity of renal cancer patients and develop an accurate and comprehensive risk model to stratify renal cancer patients for the sake of designing individualized treatment plans in terms of prognostic prediction and drug selection.

With the advancement of sequencing technology and a variety of machine learning algorithms, bio-omics has exploded, which has greatly promoted people’s understanding of tumors at the molecular level(5). At the same time, with the development of data science, a variety of robust cluster algorithms have pushed the progression of multi-omics bioinformatics science. Omics data reflect the various biological processes of cancer and provide a detailed description of the molecular mechanism of cells. Different levels of omics data reflect different relationships between the distribution of the whole genome and the occurrence, development and prognosis of cancer(6). Unfortunately, there is a lack of typing studies based on multiple omics prognostic indicators in renal cancer omics research. In this study, we performed a stratification analysis of ccRCC patients by integrating multi omics and advanced cluster algorithms and found that different subgroups presented different landscapes in each single omics data set (7-16). Finally, we used Borta algorithm to reduce the dimension of subtype specific signatures and constructed a robust risk stratification model.

## MATERIALS AND METHODS

### Data collection and preprocessing

The research workflow of this study is shown in **Figure S1**. TCGA database is a comprehensive database of the American Cancer and Tumor Project Gene Atlas Project consisting of multiple omics data of different tumor types. We downloaded ccRCC-related datasets of 530 ccRCC samples and 71 normal control samples from TCGA, which contains RNA-seq profiles, miRNA-seq profiles, the Illumina 450K DNA methylation array and WES (Whole Exome Sequencing) data. After data filtration, we finally included 258 cases of ccRCC with complete multi-omics information and clinical information into the subsequent study. But as the purpose of this study was to conduct risk stratification model through multi-omics data, we used the Cox Proportional Typing model by R Survival package to screen the signatures in each set of omics data that are closely related to the prognosis of KIRC patients for subsequent analysis, knowing that huge multi-omics data might consume too much computing resources and introducing all omics data into the study could cause the model to overfit.

### Cluster analysis

The most important parameter in any clustering study is the optimal number for cluster, where k needs to be small enough to reduce noise but on the other hand large enough to retain important information. Herein we referred to Clustering Prediction Index (CPI) and Gaps-statistics to estimate the optimal number of clusters in this study, which turned out to be 2 (12, 17). We then divided 258 ccRCC patients into two groups by clustering their multi-omics database. Ten clustering algorithms were included in the clustering process: SNF, PINSPlus, NEMO, COCA, LRAcluster, ConsensusClustering, IntNMF, CLMLR, MoCluster and iClusterBayes. Based on consensus ensembles, the results of different clustering algorithms were integrated, and the unified samples were clustered into the same category in different algorithms, namely 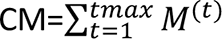 and cmij ∈[0,10] (18). Finally, we calculated the consensus matrix to represent robust pairwise similarities for samples because it considers different multi-omics integrative clustering algorithms. Then, dimensionality reduction and visualization of subtype data at multi-omics level were performed based on the idea of t-SNE graph dimensionality reduction using R ‘Rtsne’ package.

### Comparison between subgroups

Prognostic analysis of overall survival (OS) and progression-free interval (PFI) was performed based on the log-rank test between CS1 (Cancer Subtype 1) and CS2 (Cancer Subtype 2) subtypes obtained in the cluster analysis above. Differentially expressed genes (DEGs) between different subgroups were determined using the R DEseq2 package and visualized by R EnhancedVolcano package. The screening criterion for DEGs was set at adjusted *p* value<0.01 and abstract logFC=1.8. Gene ontology (GO), Kyoto encyclopedia of genes and genomes (KEGG) and Gene set enrichment analysis (GSEA) were performed to identify enriched GO terms, pathways and hallmarks by using the R package with a screening cutoff value at *p* value <0.05 and an adjusted *p* value <0.2(19). Gene set variation analysis (GSVA) was performed to identify cancer-related biological processes using R ‘GSVA’ package (20). The cancer-related hallmarks were downloaded from MSigDB database (https://www.gsea-msigdb.org/gsea/msigdb/).

### Immunity analysis and immunity-related gene expression

To quantify the proportion of immune cells in the two subtypes, several immune-related algorithms were used to calculate the cellular components or immune cell enrichment scores in the ccRCC tissue, including TIMER, CIBERSORT, QUANTISEQ, MCPCOUNTER, XCELL and EPIC (21-24). Single sample gene set enrichment analysis (ssGSEA) was employed to quantify the relative abundance of 28 immune cells in the tumor microenvironment (TME) of ccRCC (20). Differences in immune cell infiltration in TME were visualized by Heatmap and boxplot. R ‘estimate’ package was used to identify the stromal component and immune component between the two subgroups. The Tumor Immune Dysfunction and Exclusion (TIDE) (http://tide.dfci.harvard.edu) algorithms were applied to predict the immunotherapy response of each ccRCC patient.

### Copy number variation (CNV) analysis

As the main analysis tool for somatic mutation data analysis and visualization, the R ‘maftools’ package provides the possibility to compare the differences between different subtypes at the WES level(25). Through the correlation function of this package, we analyzed the tumor mutation panorama, base conversion and transversion, amino acid mutation hotspot, mutation frequency of mutation alleles, copy number mutation, mutual exclusion or coexistence mutation, and gene mutation survival in different subtypes of ccRCC patients. At the same time, the differences in drug-gene interactions and carcinogenic signaling pathways were also analyzed in this function module. Recurrent broad and focal somatic copy-number alterations (SCNAs) analysis was performed by the GISTIC 2.0 algorithm (https://www.genepattern.org/modules/docs/GISTIC_2.0) on the basis of Euclidean distance of threshold copy number at recurring peaks using Ward’s method (26).

### Drug sensitivity analysis

The response of ccRCC patients to chemotherapy drugs and small-molecule prodrugs was predicted based on the public pharmacogenomics database - Genomics of Drug Sensitivity in Cancer (GDSC; https://www.cancerrxgene.org)(27). The half maximal inhibitory concentration (IC50) was estimated by R package ‘pRRophetic’ (28). The prediction process was implemented by R package “pRRophetic”, where IC50 of the samples was estimated by ridge regression and the prediction accuracy was evaluated by 10-fold cross-validation based on the GDSC training set. Finally, the molecular structure tomographs of these candidate drugs were obtained from PubChem (https://pubchem.ncbi.nlm.nih.gov/).

### Construction of the subtype landscape

To facilitate visualization and uncover the underlying structural distribution of individual ccRCC patients, we applied the graph learning-based dimensionality reduction technique to the risk related gene expression profiles (29) in an attempt to visualize the distribution of ccRCC subtypes across individual patients using reduceDimension function of Monocle package with a Gaussian distribution. The maximum number of components was set to 4, and the discriminative dimensionality reduction with trees was used. *P*-value<0.05 was considered statistically significant.

Finally, the subtype landscape was visualized with the function plot cell trajectory with color-coded ccRCC subtypes.

### External data validation

In this part, we aimed to predict the possible subtypes of each sample in the external dataset. In most cases, this multi-classification issue is tricky for the simple reason that it is nearly impossible to entirely match recognized markers in the external cohort, so the reliability of model-based predicting methods might be low. Surprisingly, MOVICS package containing two model-free means solved this puzzle (18). First, MOVICS switches to nearest template prediction (NTP) which can be flexibly applied to cross-platform, cross-species, and multiclass predictions without any optimization of analysis parameters. Then, we compared the survival outcome of the predicted cancer subtypes in external cohort, and further checked the agreement between the predicted subtype and AJCC classification. In addition to NTP, MOVICS provides another model-free approach to predict subtypes, which first trains a partition around medoids (PAM) classifier in the discovery (training) cohort to predict the subtype for patients in the external validation (testing) cohort, and each sample in the validation cohort was assigned to a subtype label whose centroid had the highest Pearson correlation with the sample. Finally, the in-group proportion (IGP) statistic was performed to evaluate the similarity and reproducibility of the acquired subtypes between discovery and validation cohorts. In the end, we used Kappa statistics to check the consistency between different prediction results.

### Dimensionality reduction and risk model construction

Based on the multi-omics classification, we obtained two subtypes, CS1 and CS2. In the previous analysis, the DEGs between the subtypes were obtained and the DEGs that were positively correlated with the signals of the CS1 and CS2 subtypes were divided into risk score signature A and B respectively(30). Furthermore, the dimension reduction of the risk gene signatures A and B was conducted using Boruta algorithm, and Principle Component Analysis (PCA) algorithm was employed to draw principal component 1 as the score(31). Finally, we ran a previously reported method to get the risk score of each patient: score=∑ *PC*1*A*-∑*PC*1*B*. According to the median value of the patient’s risk score, the patients were divided into a high-risk group and a low-risk group, and survival differences between the two groups was analyzed using R ‘survival’ package.

### Statistical analysis

Spearman or Pearson correlation analysis was used to calculate the correlation coefficient between two variables. Wilcoxon rank-sum or T test was applied to evaluate the difference when comparing between two continuous variables. Kruskal-Wallis or ANOVA test was introduced to compare difference among three or more groups. Also, chi-squared or Fisher’s test was applied to compare difference in categorical variables. The nomogram was developed to assess individual outcome of ccRCC patients by using rms package. Calibration curves were calculated by calibrate function implemented in rms package. Univariate Cox regression and multiple Cox regression analyses were performed to calculate the hazard ratios (HRs). The receiver operating characteristic (ROC) curves were plotted by ‘timeROC’ R package. Area under the ROC curve (AUC) and Harrell’s concordance index (C-index) were employed to evaluate the performance of the risk score in predicting OS and PFI. All operations are in R 4.0.3 (https://www.r-project.org). P values were two-sided and P-value < 0.05 was considered statistically significant. If multiple comparisons were involved, p-values were adjusted using Benjamini–Hochberg method. Visualization was achieved by R ggplot2, ggpubr and ComplexHeatmap packages.

## Results

### Multi-omics landscapes of the two ccRCC subgroups

The number of data clusters was estimated using CPI and Gap statistics in combination with the aforementioned 10 algorithms. Both the correlation matrix heatmap and the silhouette map showed good inter-group heterogeneity and intra-group consistency between the two subgroups, indicating that our classification not only preserved the heterogeneity of different types of ccRCC but reduced unnecessary redundancy **Figure S2**. The two types of ccRCC subtypes presented different landscapes in different omics including mRNA, lncRNA, miRNA, methylation and somatic mutation levels. REG1A, KRT19, SLPI, SLC5A1, SLC22A6, CYP4A11, SLC22A12, ALDOB, FOSB and REN were miRNAs that showed significant differences between the two subgroups; RP4-764022.1, AC079630.2, AC147651.5, RP11-397G17.1, AP000439. 3, LINC00671, RP11-133F8.2, RP11-14C10.5 and LINC01426 were differential lncRNAs; has-let-7f-1, has-let-7f-2, has-mir-126,has-mir-139, has-mir-1468, has-mir-138-1, has-mir-138-2,has-mir-1269a, has-mir-1293 and has-mir-135b were differential miRNAs; cg01413054, cg23557926, cg07912922, cg06879394, cg05232694, cg19836199, cg08443845, cg04915566, cg05514299 and cg07388969 were differential methylation sites; VHL, PBRM1, KDM5C, MTOR, TTN, BAP1, MUC16, HMCN1, ATM and SETD2 were differential genes related to CNV (**Figure 1A**). The differences between the two tumor types were observed in the t-distributed stochastic neighbor embedding (t-SNE) dimensionality reduction distribution maps of each omics (**Figure 1B**). Prognostic analysis showed that the CS2 subtype was better than the CS1 subtype in terms of OS and PFI (**Figure 1C**). Finally, the classification of this study maintained a certain degree of consistency with the traditional AJCC classification and TNM stage classification, and was able to provide more accurate classification information for ccRCC patients (**Figure 1D** **and Table 1**).

**Figure 1.**
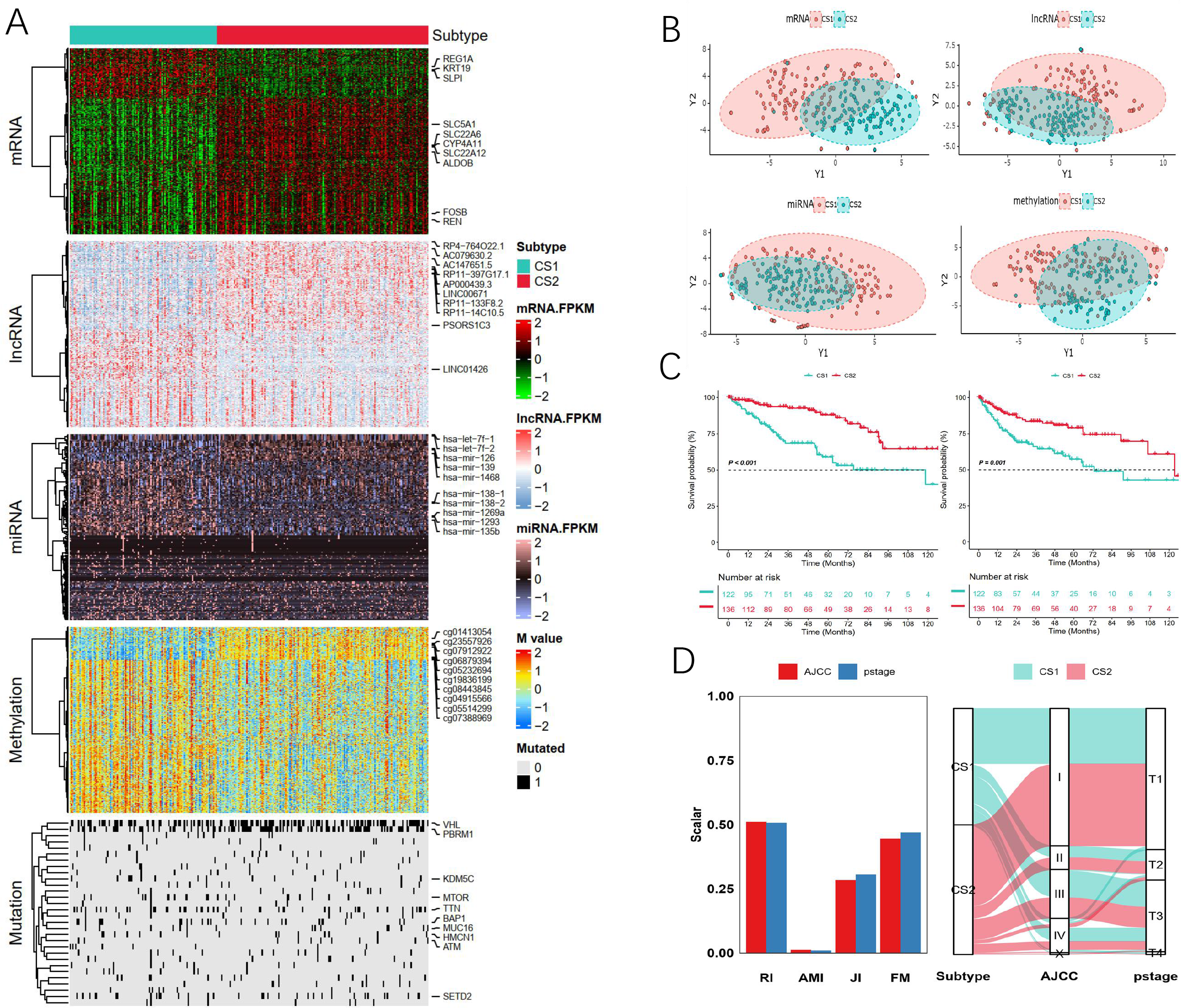
The Landscape of mutli-omics differences in ccRCC. **A.** Comprehensive heatmap of multi-omics integrative clustering with dendrogram for samples. **B.** Dot plot for two distinct clusters identified by t-SNE algorithm based on mRNA, lncRNA, miRNA and methylation profiles. **C.** Kaplan-Meier survival curve of OS and PFI of the two identified subtypes of renal cancer in TCGA-KIRC cohort. **D.** Agreement of the two identified subtypes of renal cancer with AJCC classification and pathological stage in TCGA-KIRC cohort. OS, overall survival; PFI, progression-free interval

**Table 1.**
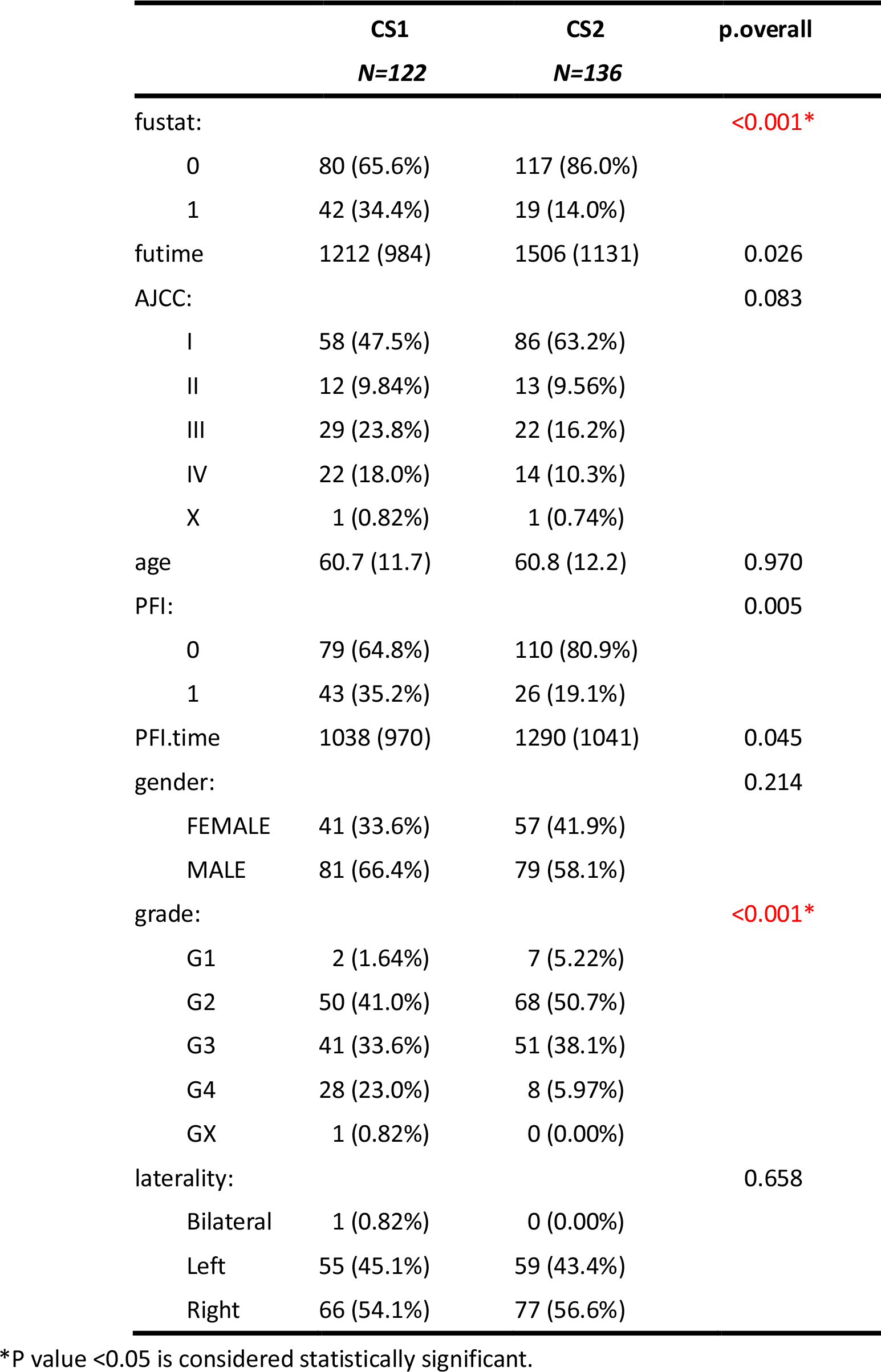
Summary descriptives table by groups of ‘cluster’

### Pseudo-timing analysis

This analysis cast individual patients into a manifold with sparse tree structures and defined the risk landscape of ccRCC. Consistent with our previously defined subtypes, many patients were segregated into two distinct clusters and presented different states (**Figure 2A**). The location of individual patients in the risk landscape represents the overall characteristics of the tumor immune microenvironment in the corresponding subtype. For instance, the high-risk subtype CS1 and low-risk subtype CS2 were distributed at the opposite end of the horizontal axis in the risk landscape (**Figure 2B**). Therefore, we hypothesized that the horizontal axis in the risk landscape represented the overall risk score. However, the vertical coordinate of the developmental trajectory appeared to be more complex and may reflect several characteristics. The risk landscape further revealed significant intra-cluster heterogeneity within each subtype. We observed that certain subtypes appeared to be more diverse and heterogeneous than others. For instance, CS1 could be further divided into three subgroups based on their location in the risk landscape, which showed different risk gene expression profiles in specific modules (**Figure 2C**). Interestingly, the three subgroups of patients in CS2 as stratified by the risk landscape were associated with distinct prognoses (**Figure 2E-F**). Similar results were obtained for CS2 (**Figure 2D**). The result of prognosis analysis showed two distinct subtypes in the risk landscape within CS2, in which 2B showed poorer prognosis than other types in terms of OS and PFI (**Figure 2G-H**). These results indicate that our risk landscape analysis can provide a complementary value to previously identified risk subtypes.

**Figure 2.**
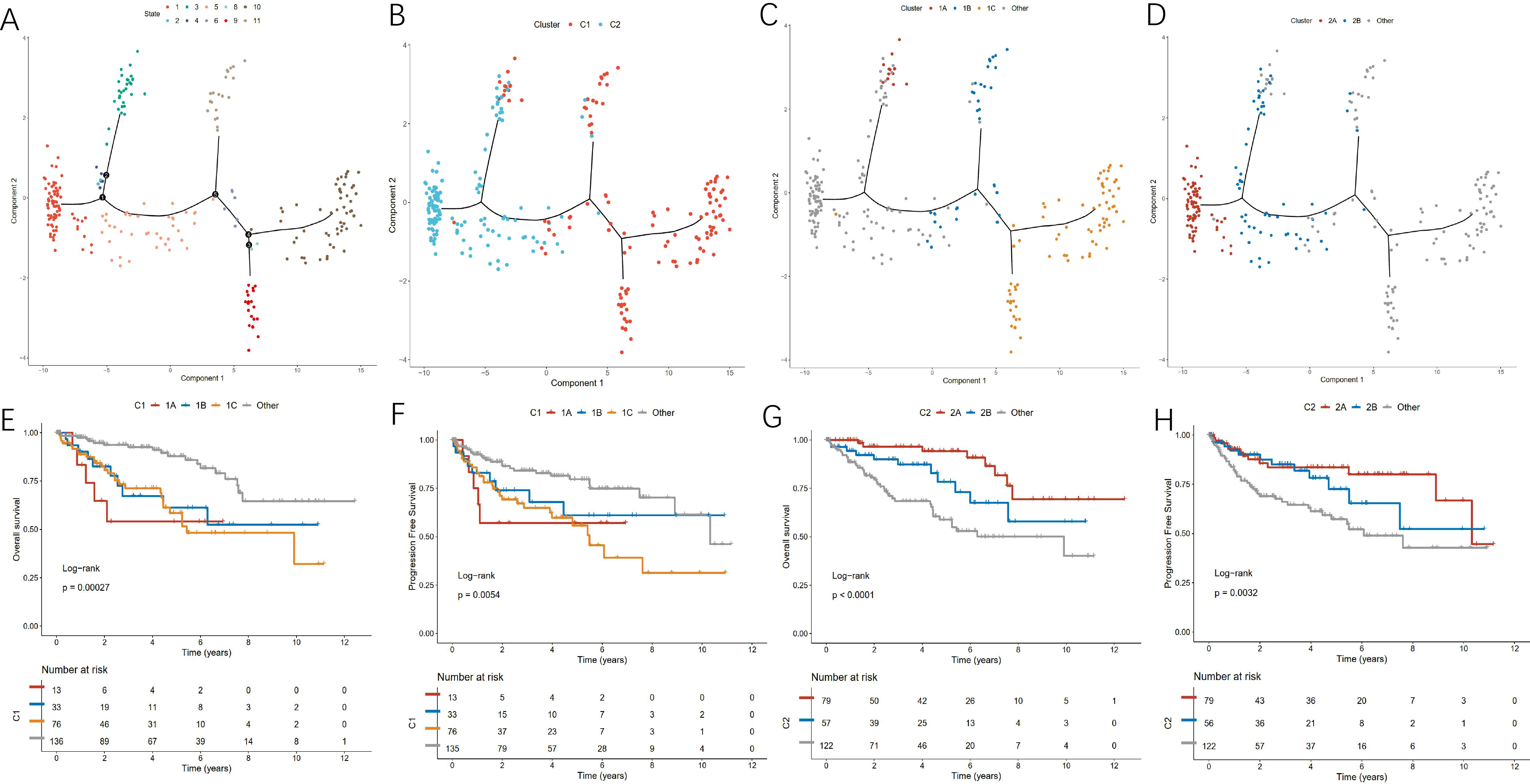
The risk landscape of ccRCC and the intracluster heterogeneity within subtypes. **A.** The subtype of ccRCC clustered by state. **B.** The risk related mRNA landscape of ccRCC: each point represents a patient with colors corresponding to the subtype defined previously. **C.** Patients of CS1 subtype could be further stratified into three subgroups based on their location in the risk related mRNA landscape. **D.** Patients of CS2 subtype could be further stratified into two subgroups based on their location in the risk related mRNA landscape. **E-F**. The three subgroups of patients in CS1 as stratified by the risk related mRNA landscape were associated with distinct prognoses of OS and PFI. Log-rank P value was calculated among subgroup stratification. **G-H**. The two subgroups of patients in CS2 as stratified by the risk related mRNA landscape were associated with distinct prognoses of OS and PFI. Log-rank P value was calculated among subgroup stratification.

### Functional enrichment analysis

Through differential analysis, we identified a total of 950 DEGs, which are shown in volcano plot (**Figure 3A**, **Figure S3**), including 907 up-regulated genes and 43 down-regulated genes, and biomarker signatures of each subtype are provided in **Supplement Table 1**. GO analysis showed that the differential genes between CS1 and CS2 subgroups were enriched in cornification, epidermis development, keratinocyte differentiation, skin development, negative regulation of endopeptidase activity in Biological Process (BP) module. In Cellular Compartment (CC) module, DEGs were enriched in the collagen-containing extracellular matrix, intermediate filament, intermediate filament cytoskeleton, high-density lipoprotein particle, plasma lipoprotein particle, as well as in peptidase inhibitor, endopeptidase regulator, endopeptidase inhibitor, serine-type endopeptidase inhibitor and peptidase regulator activities (**Figure 3B**). KEGG pathway analysis revealed differential pathways such as complement and coagulation cascades, while molecular function (MF) analysis revealed EMC-receptor interaction and p53 signaling pathway (**Figure 3C**). Through GSEA analysis, we found that extracellular matrix organization, formation of the cornified envelope, keratinization, SLC-mediated transmembrane transport and transport of small molecules, were significantly enriched in CS1 subgroup (**Figure 3D**). GSVA analysis showed that hypoxia, peroxisome, KRAS signaling pathway, myogenesis and coagulation were significantly enriched in CS1, while G2M checkpoint, fatty acid metabolism, heme metabolism and mitotic spindle were significantly enriched in CS2 (**Figure 3E**).

**Figure 3.**
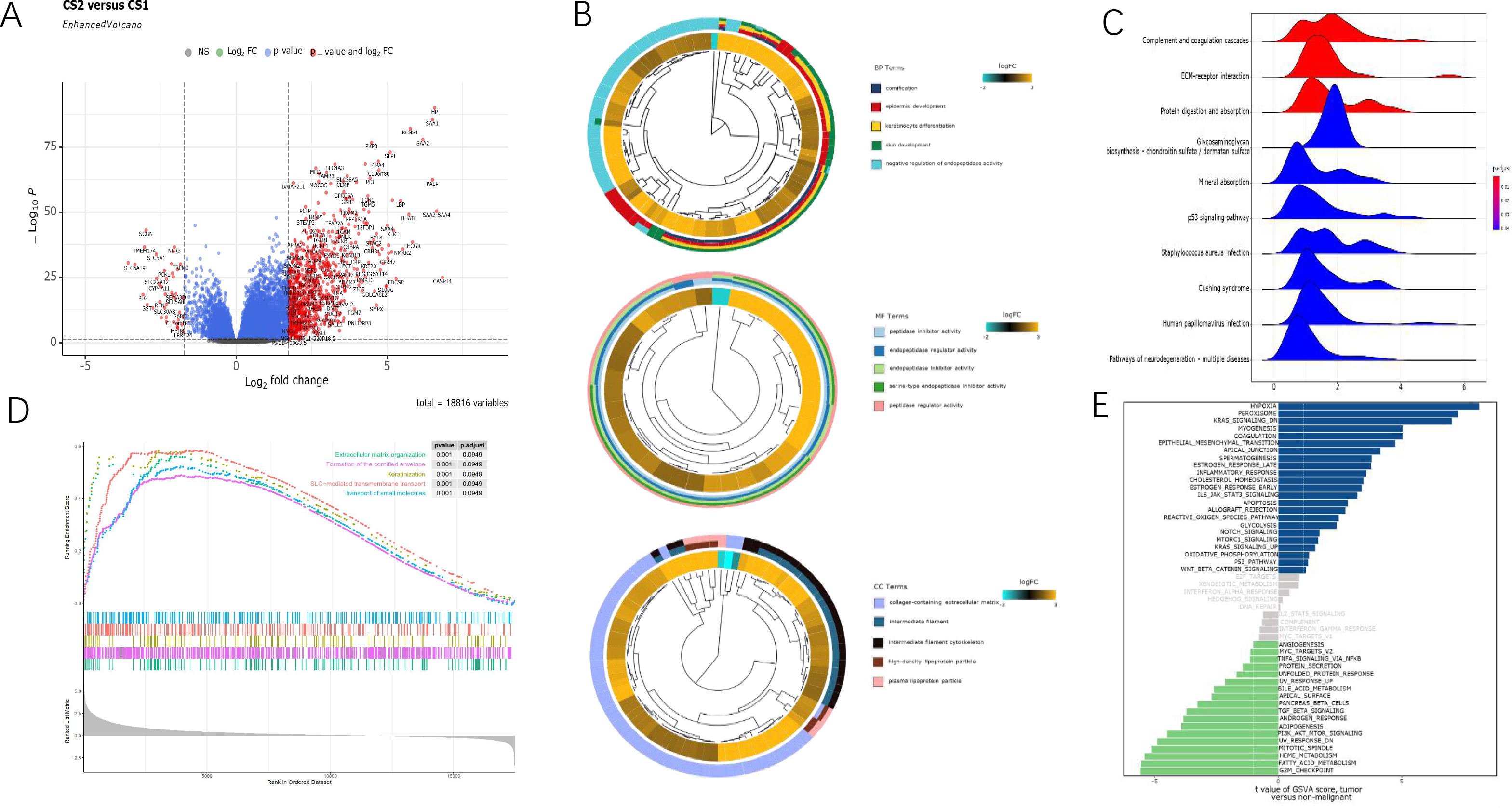
Functional enrichment analysis of DEGs between CS1 and CS2 subtypes. **A.** Volcano map of different expression genes. **B.** GO enrichment analysis. **C.** KEGG pathway analysis. **D.** Gene set enrichment analysis (GSEA): hallmarks in the subgroups. **E.** GSVA: differences in enrichment between the subgroups.

### Comparison of somatic mutations

We had previously analyzed the differences between patients with different subtypes at the transcriptome level, and in the present study we tried to analyze the differences between the subtypes at the genome level, and further determine whether there was evidence of the disparity between CS1 and CS2 subgroups of patients at the genomic level. It was found that CS1 and CS2 presented different mutation landscapes. The top 20 most common mutation genes are shown in **Figure 4A**. Among them, the mutation rate of VHL, PBRM1 and ARID1A in CS1 subgroup was significantly lower than that in CS2 group (41% *vs.* 54%; 26% *vs.* 54%; 1% *vs.* 7%), while the mutation rate of TTN in CS1 subgroup was significantly higher than that in CS2 subgroup (15% *vs.* 21%; 4% *vs.* 17%; 10% *vs.* 16%) (**Figure 4B**). To be noted, PBRM1 and BAP1 were the most significantly different mutant genes between the two subgroups (**Figure S4A**). Some previous studies (32, 33) identified VHL, PBRM1 and TTN as the top three mutations in the two subtypes of patients, and they were involved in the occurrence and progression of ccRCC. However, it was found in the present study that the mutation frequencies of the above genes were not consistent between the two subgroups of patients, suggesting that the classic ccRCC mutations play totally different roles in the CS1 and CS2 subtypes of ccRCC. For instance, PBRM1 is the second largest mutation gene in renal cancer, while the mutation event of PBRM1 in the two renal cancer subtypes described herein presented inconsistent survival benefits for the patients. PBRM1 mutation was identified as a protective factor in CS2 subgroup (**Figure 4C**). Similarly, the mutation sites were not consistent between the two subgroups of ccRCC (**Figure 4B**). Next, we investigated the co-occurring and exclusive mutations of the top 25 most frequently mutated genes using CoMEt algorithm, and found that there existed some specific co-mutations in CS1 subtype such as PBRM1-VHL, PBRM1-NOTCH2 and PBRM1-KMT2C, and also there were specific co-mutations such as ARID1A-ATRX5, SETD2-THSD7B, ARID1A-DNAH9 in CS2 group, suggesting that they probably played a redundant role in the same pathway and harbored the selective advantages between them to keep more than one copy of the mutations (**Figure 4D**). However, the ratio between transversion (Tv) and transition (Ti) in all SNVs was approximately 1:1 and remained stable in both cohorts (**Figure S4B**). The distribution of variant allele frequency (VAF) in different genes was found to be inconsistent between the two subtypes. Among them, the genes with the highest VAF in CS1 group were KDM5C, VHL and PBRM1, while the first three VAF genes in CS2 group were VHL, PBRM1 and ATM (**Figure S4C**). Finally, there were differences in the sudden change landscape between the two subgroups. The different gene mutations showed distinct clinical outcomes for patients in CS1 and CS2 subgroups (**Figure S3D**). For example, PBRM1 was a typical example to demonstrate the different mutation spots between the two cohorts and the plausible chain reaction of the differences took place in prognostic impact (**Figure S4E**). At the same time, based on the differences in CNV data between the two subgroups, we explored possible treatments based on the mutation data of the two subtypes using the drug-Interactions function in the maftools package and the DGIdb database (https://www.dgidb.org/). Among them, the possible therapeutic targets in CS1 group mainly included BAP1, COL24A1, DST, HMCN1 and KDM5C. The potential therapeutic targets in CS2 group mainly included ATM, COL6A6, DST, ERBB4 and HMCN1 (**Figure 4E**).

**Figure 4.**
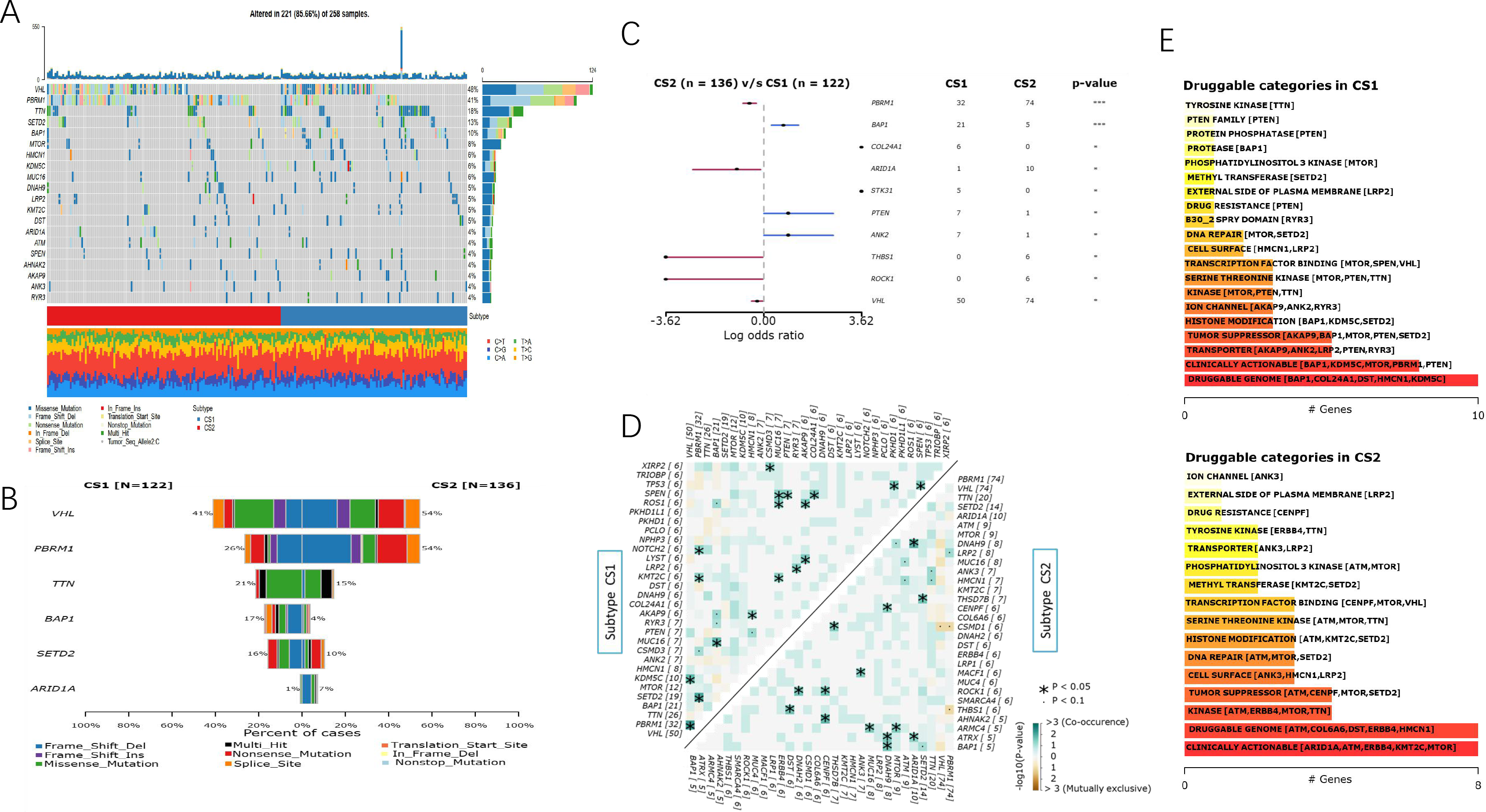
Landscape of Somatic Mutation in CS1 and CS2 subgroups. **A-B.** Waterfall plot shows the mutation distribution of the top 20 most frequently mutated genes. The central panel shows the type of mutation in each ccRCC sample. The upper panel shows the mutation frequency of each ccRCC sample. The bar plots on the left show the frequency and mutation type of genes mutated in 258 ccRCC samples. The lower part shows different subgroups including CS1 and CS2. **C.** Forest plot displays the top 6 most significantly differentially mutated genes between the two subgroups. **D.** Heatmap illustrates the mutually co-occurring and exclusive mutations of the top 25 frequently mutated genes. The color and symbol in each cell represent the statistical significance of the exclusivity or co-occurrence for each pair of genes. **E.** Potential druggable gene categories from mutation dataset in CS1 and CS2.

### CNV differences between CS1 and CS2 subgroups of ccRCC

CNV differences were compared between patients with different subtypes of KIRC. The incidence of the rate of copy number variation in CS2 was higher than that in CS1 (**Figure 5A**). GISTIC2.0 was used to define recurrently amplified and deleted regions in the two subgroups(**Supplement Table 2**). The results showed that the two subtypes had frequent CNVs in the region containing oncogenes, tumor suppressor genes (e.g. VHL and PBRM1) and metabolism regulator (e.g. COL4A3 and COL4A4), indicating that CNVs might play a significant role in the tumorigenesis and progression of ccRCC. We found recurrent focal CNVs in CS1 including amplifications containing 5q35.1(ADRA1B), 3q26.33 (PIK3CA) and deletion of 9p21.3(CDKN2A) and 3p21.31(KIF9). Recurring focal CNVs in CS2 included amplifications of 5q33.2(KIF4B), and deletion of 3p11.1(HTR1F). These specific CNVs might contribute to the formation of the two subtypes (**Figure 5B**).

**Figure 5.**
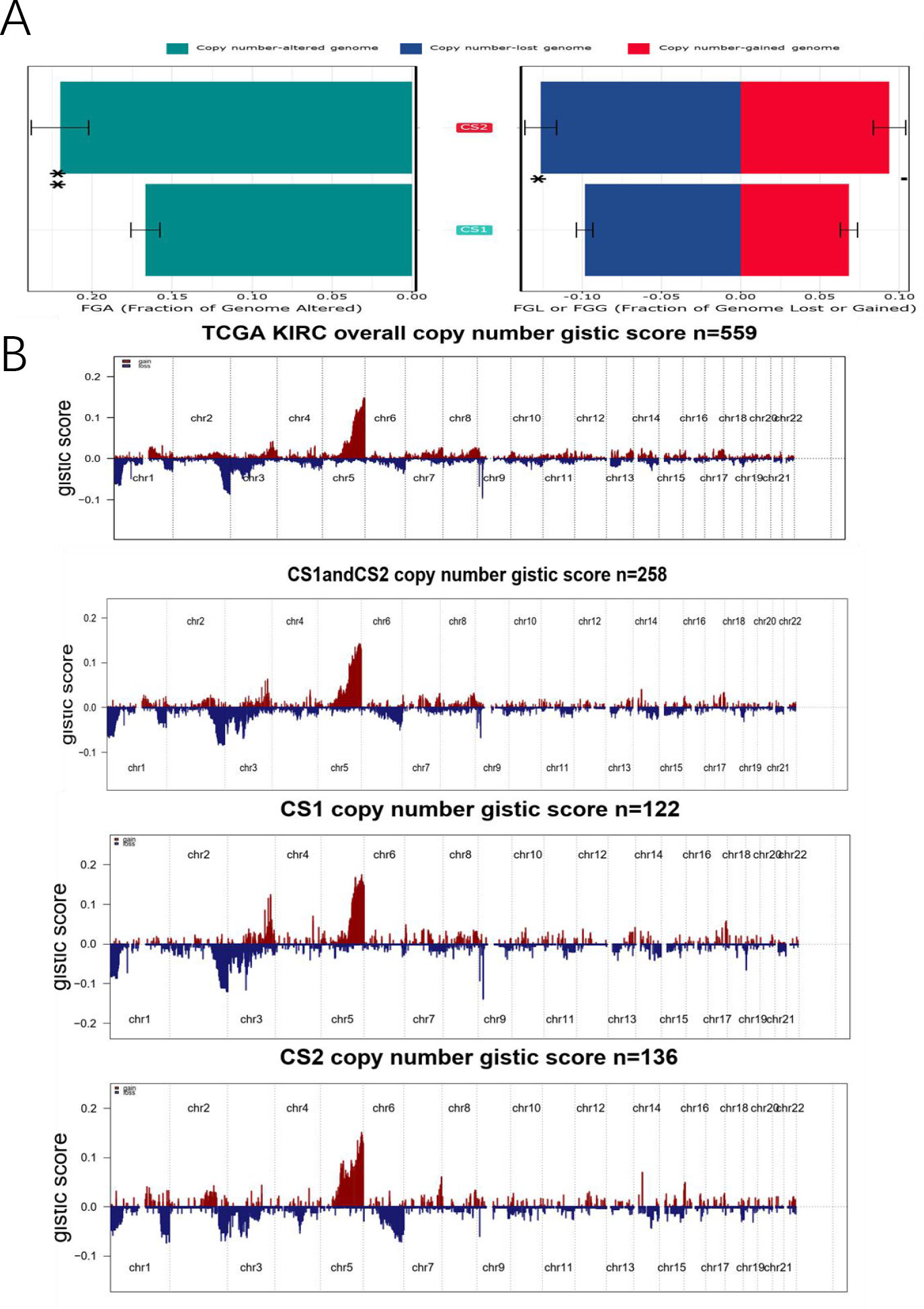
CNV differences between subgroups. **A.** Barplot of fraction genome altered in the two identified subtypes of renal cancer in TCGA-KIRC cohort. **B.** Composite copy number profiles for ccRCC with gains in red and losses in blue and grey highlighting differences between CS1 and CS2.

### Carcinogenic pathway differences between CS1 and CS2 subtypes of ccRCC

Through CNV data, we found that onco-pathways were not completely consistent between the two subgroups. Among them, the most critical carcinogenic pathways in CS1 subgroup were RTK-Ras, Notch, PI3K, Hippo, WNT, TP53, MYC, NRF2, and Cell Cycle *vs.* RTK-Ras, Notch, Hippo, WNT, PI3K, TP53, MYC, Cell Cycle, NRF2, TGF_β and other pathways in CS2 subgroup (**Figure 6A**). At the same time, GSVA analysis was exploited to verify the difference in onco-pathways between the two subgroups (**Figure 6B**).

**Figure 6.**
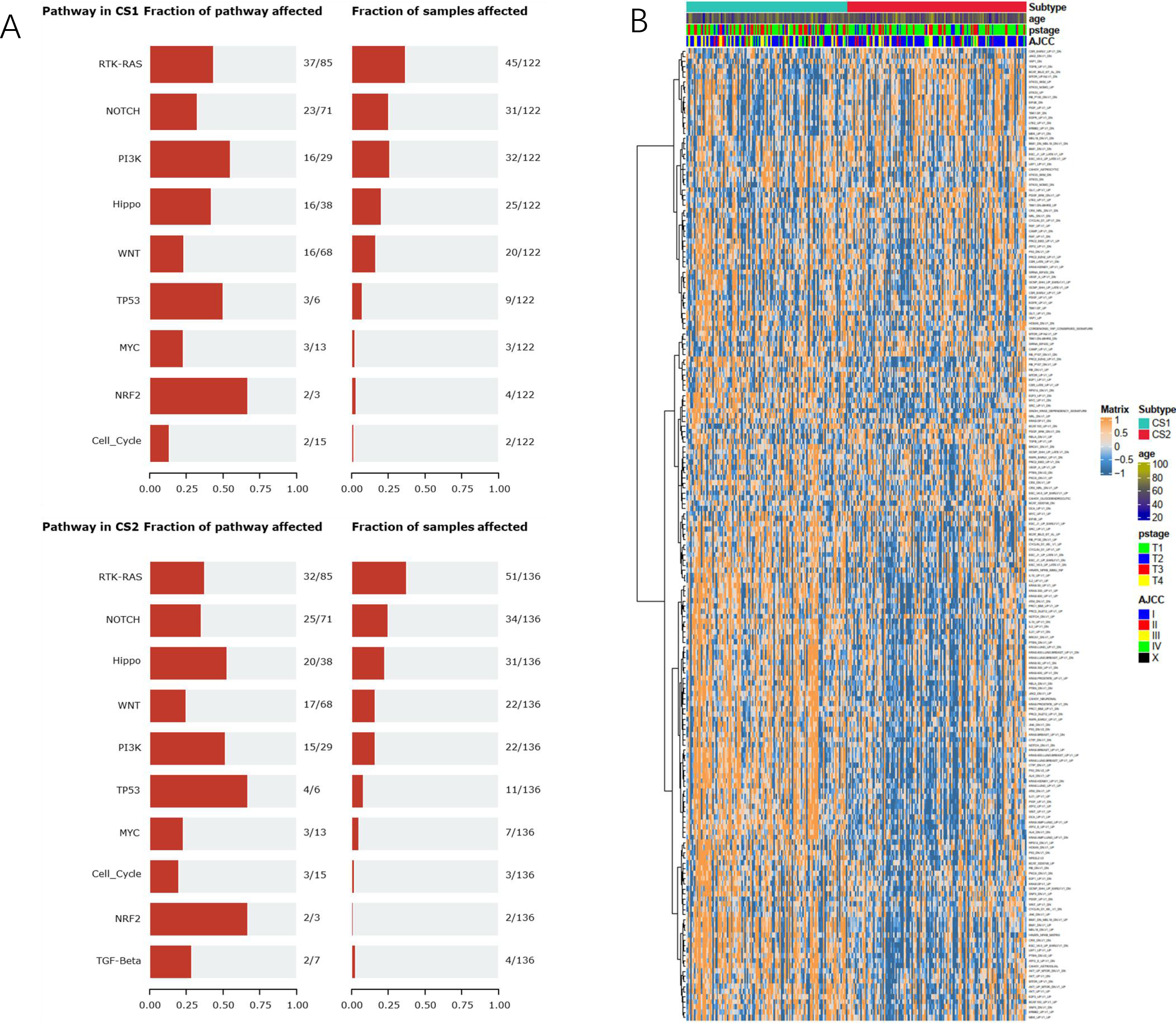
Carcinogenic pathway differences. **A.** Differences in oncogenic signaling pathways between CS1 and CS2 subgroups. **B.** Heatmap of oncogenic pathways based on ssGSEA algorithm in CS1 and CS2 subgroups.

### Immune infiltration analysis

The result of our immune infiltration analysis showed a significant difference in the immune landscape between CS1 and CS2 subgroups. The algorithms showed fewer infiltrating cells in CS2 subgroup as compared with CS1 subgroup (**Figure 7A**), and the expression of marker genes related to the immune microenvironment was also significantly different between the two groups (**Figure S5**). Among them, the chemokine family, costimulatory factors, co-inhibitory molecules, interferon and MHC molecules were included in CS1 subgroup. GSVA analysis showed that Differences in specific immune cell infiltration between the two groups were calculated and the result showed a higher degree of multiple immune cell infiltration in CS1 group than that in CS2 group (**Figure 7B**). GSVA analysis showed that significantly more immunity-related pathways were activated in CS1 subtype as compared with CS2 subtype, including interleukins, cytokines, B cell functions and T cell functions (**Figure 7C**). However, there was no significant difference in tumor mutation burden (TMB) between the two subgroups (**Figure 7D**). However, TIDE algorithm showed a significantly higher immune check inhibitor response rate in CS2 patients compared with CS1 patients (*p*<0.01) (**Figure 7E**). Finally, with respect to differentially expressed common immune reverse regulator makers between the two groups, the expression of PD1 in CS2 subgroup was higher than that in subgroup (*p*=0.083), while PDL1 in CS2 subgroup was significantly lower than that in CS1 subgroup (*p*<0.01). But other common immunosuppressive markers including TIGIT, LAG3, CTLA4 and KLRB1 were not significantly different between the two subgroups (**Figure 7F**). The ESTIMATE results showed that the Immunescores and Estimatescores in CS1 group were lower than those in CS2 group (**Figure 7G**).

**Figure 7.**
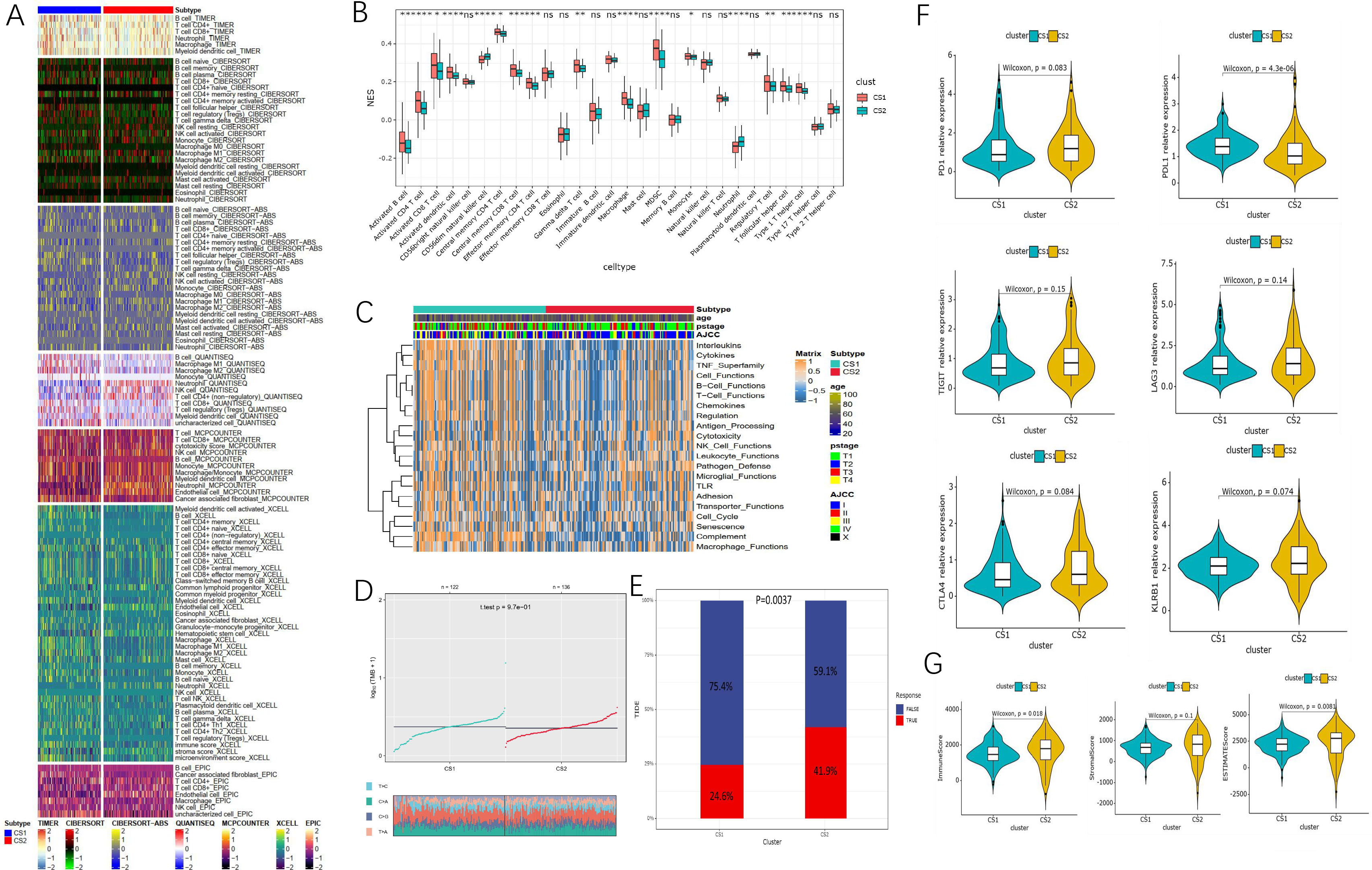
Identification of immune status. **A.** Heatmap of tumor-related infiltrating immune cells based on TIMER, CIBERSORT, CIBERSORT-ABS, QUANTISEQ, MCPcounter, XCELL and EPIC algorithms in CS1 and CS2 subgroups. **B.** Different normalized enrichment scores of immune cells between the subgroups. **C.** Heatmap of different immune-related pathway enrichment scores between the subgroups. **D.** Difference in TMB between CS1 and CS2 subgroups. **E.** Difference in response of immune check point inhibitor treatment based on TIDE algorithm **F.** Expression of immune checkpoints in CS1 and CS2 subgroups. **G.** Difference in ESTIMATE scores between the subgroups

### Drug sensitivity analysis

In this section, we compared the sensitivity to Saracatinib, Lisitinib, Imatinib, Gefitinib, Erlotinib, Dasatinib, Crizobinib and Sunitinib between the two subgroups of ccRCC patients, and found no significant difference in sensitivity to Sunitinib and Erlotinib between CS1 and CS2 subgroups, but patients in CS2 subgroup were more sensitive to Sorafenib, Lisitinib, Gefitinib and Dasatinib than those in CS1 subgroup (**Figure8A**). The previous prognosis analysis showed that the prognosis of CS1 subgroup was poorer than that of CS2 subgroup. Therefore, based on the GDSC database, we explored whether there were any other sensitive drugs for CS1 subgroup, and finally discovered 53 kinds of small-molecule drugs that could be used as potential drugs for the treatment of CS1 patients (**Supplement Table3**). The top 10 molecular drugs with the most significant differences were: CGP.082996, CMK, JNJ.26854164, ZM.447439, GNF.2, LAM. A13, RO.3306, WO2009093972, CGP.60474, and VX.680 (**Figure 8B**). The detailed structures of the above molecules are provided in **Figure S6**.

**Figure 8.**
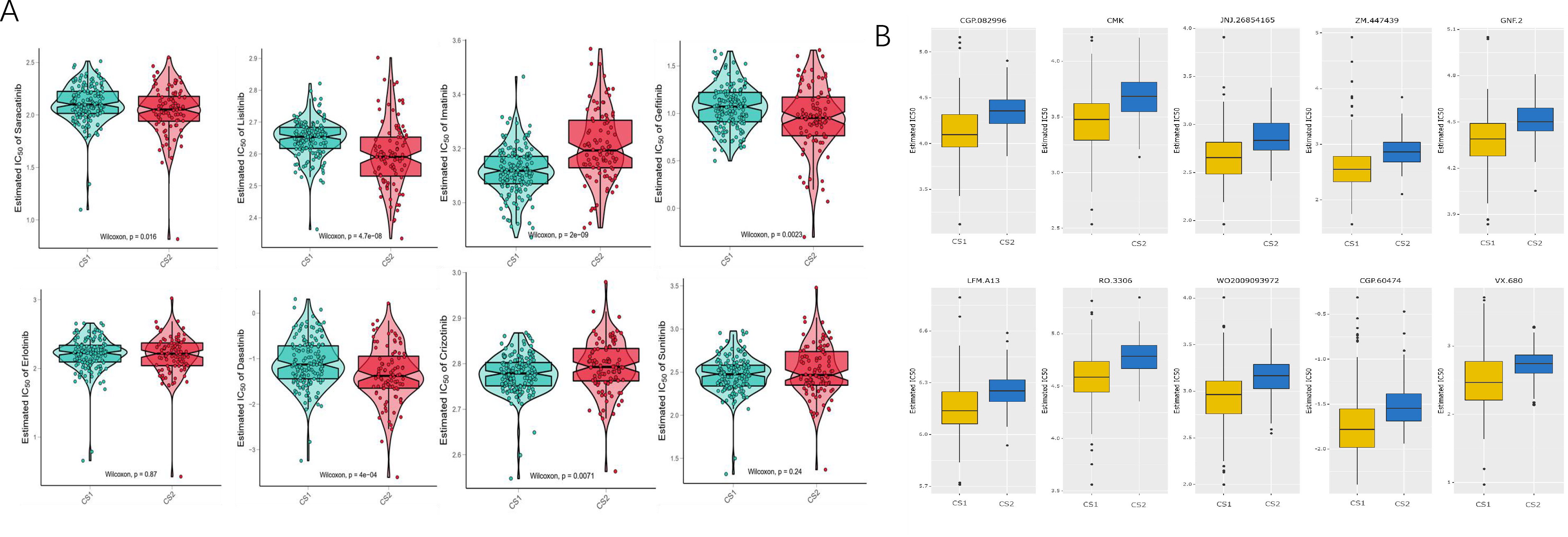
Difference of drug sensitivity. **A.** Differences in estimated IC50 of the molecular targeted drugs (Saracatinib, Lisitinib, Imatinib, Gefitinib, Erlotinib, Dasatinib, Crizobinib and Sunitinib) between CS1 and CS2 subgroups. **B.** Differences in response to 10 chemotherapy drugs between the two prognostic subtypes.

### External data verification for the subtyping method of ccRCC

To verify whether the subtype-specific signature could accurately classify renal cancer into CS1 and CS2 subtypes in an external data set, we used the remaining 273 renal cancer data set in TCGA as a validation set to evaluate the robustness of the subtype signature classier. It was found that the subtype signature could accurately verify the outer data into two subgroups (**Figure 9A**). The classification of the outer dataset was consistent with the classic AJCC classification in some extent (**Figure 9B**). Among the validation data sets, OS and PFI in CS2 group were poorer than those in CS1 group (*p* <0.01) (**Figure 9C**). Finally, we used Kappa statistics for consistency evaluation, and we compared the consistency between NTP-predicted subtype and COMIC in discovery TCGA-KIRC (scores=0.743, *p*<0.001), between PAM-predicted subtype and COMIC in discovery TCGA-KIRC (scores=0.899, *p*<0.001), and between NTP and PAM-predicted subtype in validation ICGC-KIRC (scores=0.75, p<0.001) (**Figure 9D**). The results demonstrated that our typing research was robust and reliable.

**Figure 9.**
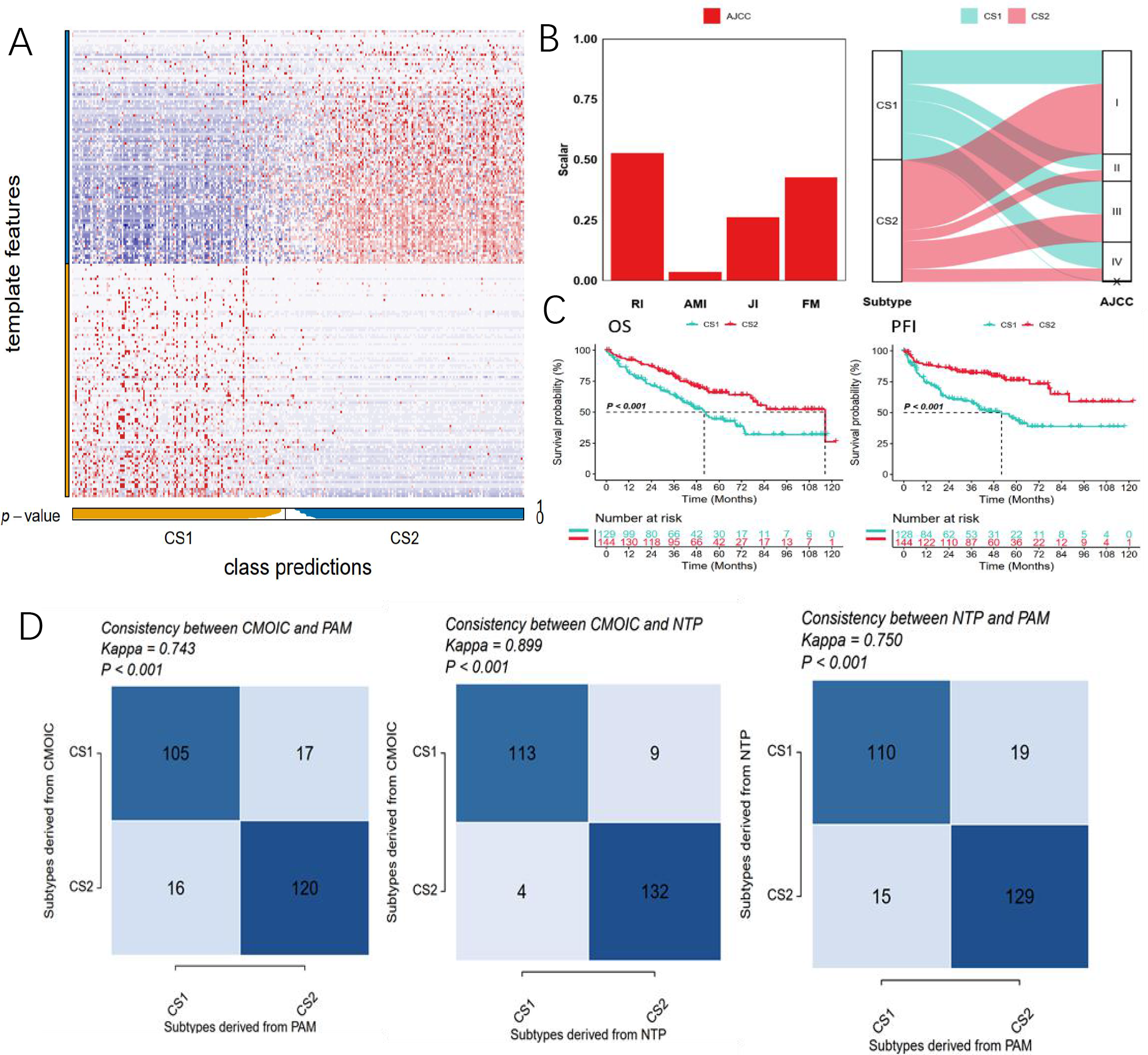
Verification in external dataset. **A.** Heatmap of NTP in out KIRC cohort using subtype-specific upregulated biomarkers identified from TCGA-KIRC cohort. **B.** Agreement of the two predicted subtypes of renal cancer with AJCC classification in outer KIRC cohort. **C.** Kaplan-Meier survival curve of the two predicted subtypes of renal cancer in outer KIRC cohort. **D.** Consistency heatmap using Kappa statistics.

### Dimensionality reduction clustering and risk model

To clarify the subtype signatures of the CS1 and CS2 subtypes, we used the DESeq2 package to perform differential analysis to determine the transcriptome differences between these subtypes, and then we performed unsupervised clustering based on the differential genes, and set the genes that were positively related to CS1 as signature B, and the genes positively related to CS2 as signature A (**Figure 10A**). Next, to reduce noise or redundant genes, we used Boruta algorithm to reduce the dimensionality of the signature genes of CS1 and CS2 subtypes (**Figure S7A**). At the same time, we chose the most critical signatures according to the degree of contribution (**Figure S7B**). Furthermore, a method similar to Gene expression grade index was applied to define the risk score of each patient(34). Based on the risk scores obtained, the patients were divided into a high-risk group and a low-risk group (**Figure 10** **B, C**). Prognosis analysis showed that the prognosis of the patients in the high-risk group was significantly poorer than that of the patients in the low-risk group in terms of OS and PFI (*p*<0.01), and the ROC curve showed that the risk score could more accurately assess the long-term survival rate of the patients (**Figure 10** **D-K**). To develop a clinically relevant quantitative method for predicting the probability of patient mortality, we constructed a nomogram by integrating the risk score and other clinical prognostic factors (**Figure 10L**). The calibration plot indicated that the derived nomogram had better performance than that of an ideal model (**Figure 10M**).

**Figure 10.**
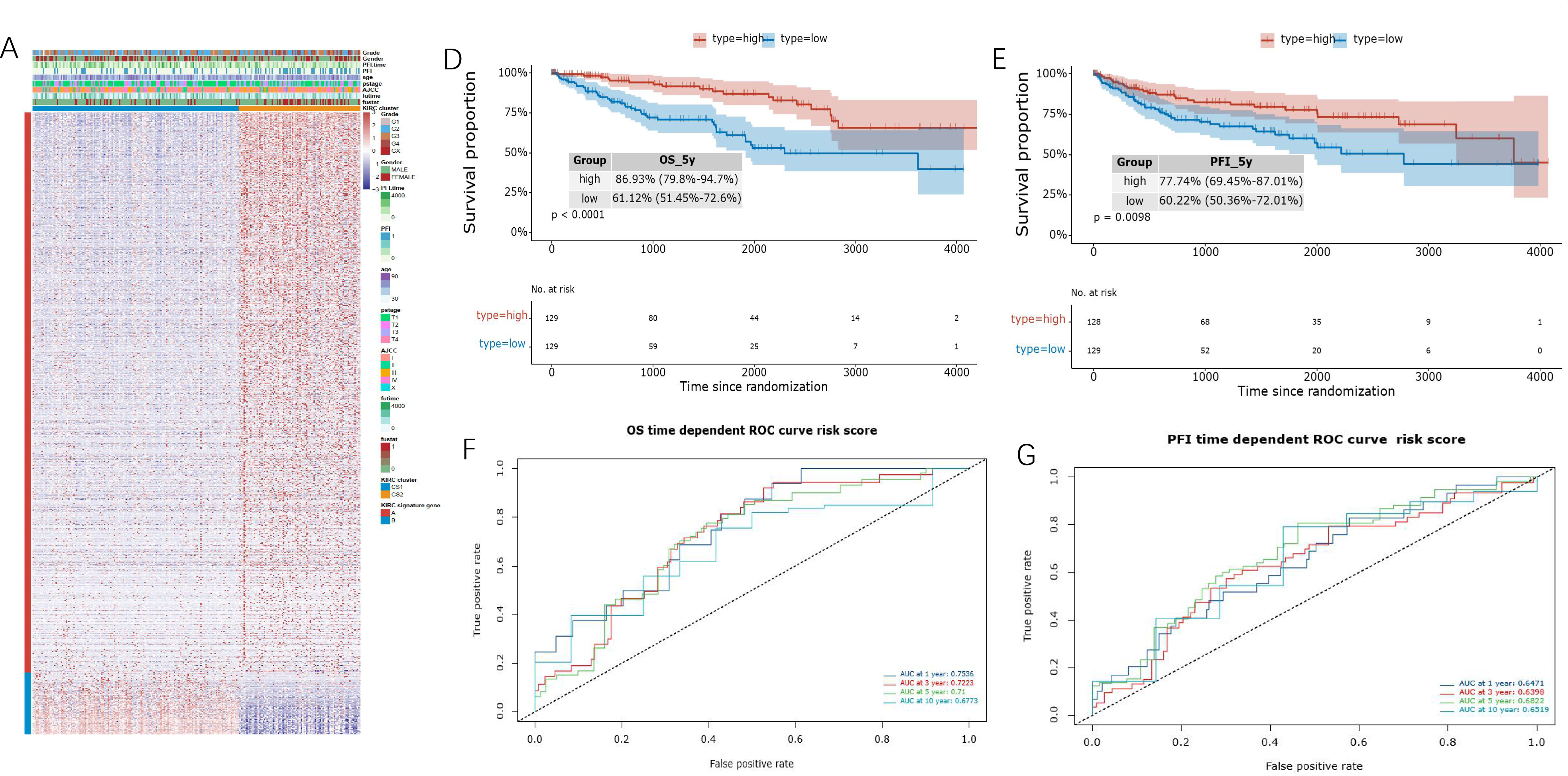
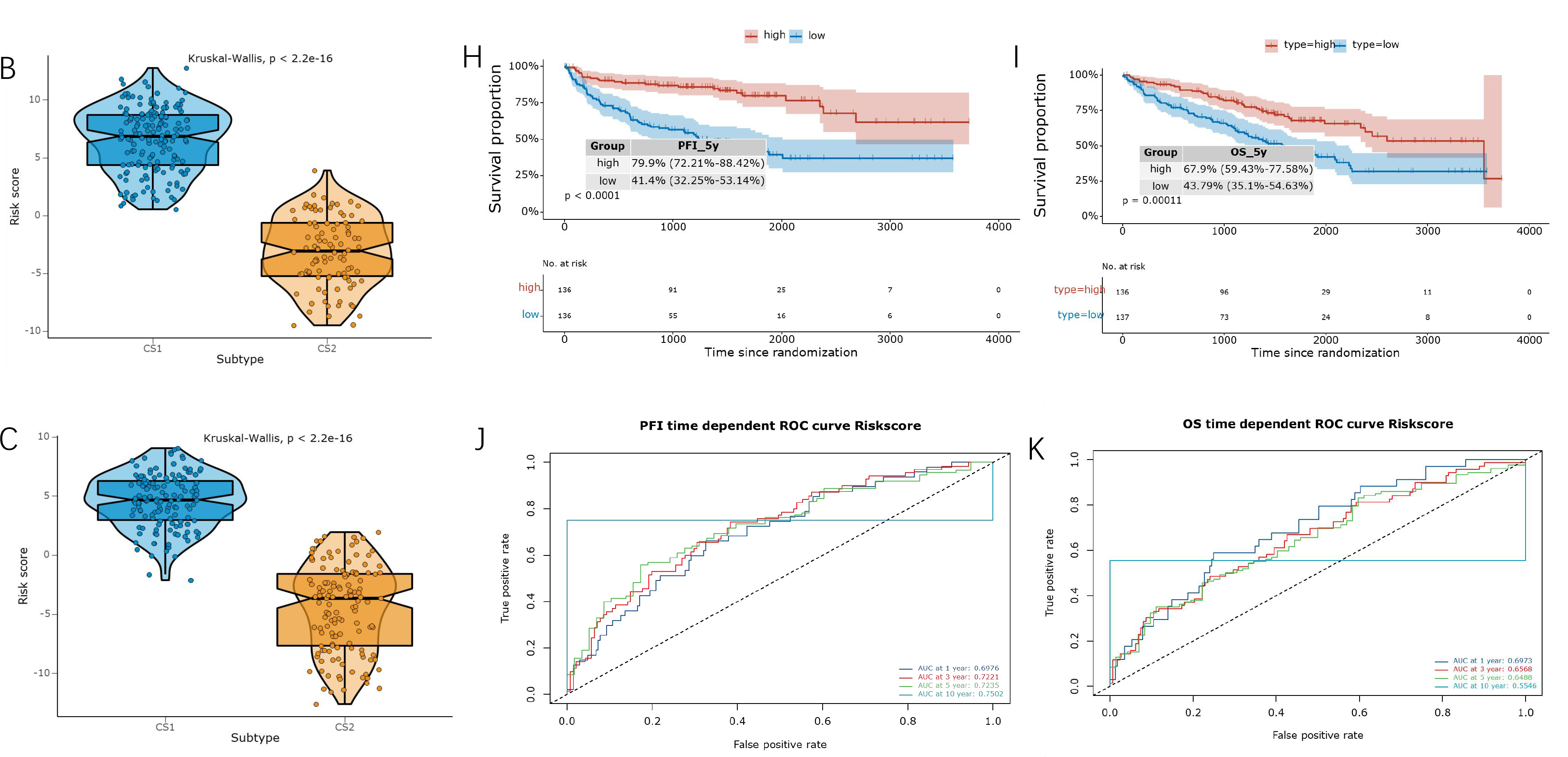
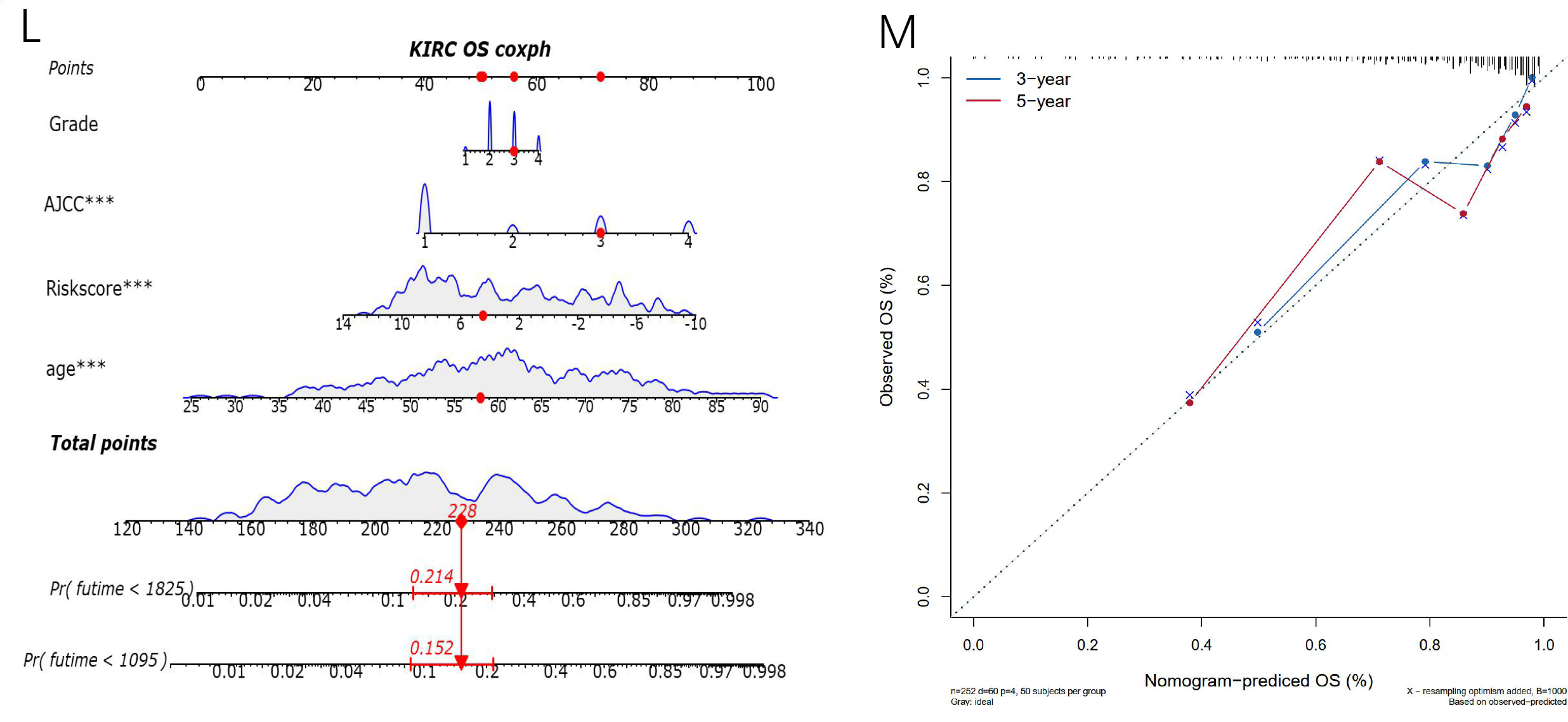
Construction of the risk scores. **A.** Common DEGs in CS1 and CS2 subtypes were classified into gene signature A and B subtypes by unsupervised clustering. **B-C**. Estimated risk score between train and validation cohorts. **D-G.** ROC curve of OS and PFI, as well as Kaplan-Meier survival curve of OS and PFI of the two identified subtypes of renal cancer in train cohort. **H-K.** ROC curve of OS and PFI, as well as Kaplan-Meier survival curve of OS and PFI of the two identified subtypes of renal cancer in train cohort. **L.** Nomograms for predicting the probability of patient mortality based on risk score of 3- and 5-year OS. **M.** Plots depict the calibration of nomograms based on risk score of 3- and 5-year OS.

## Discussion

ccRCC is significantly different from many other solid tumors. Firstly, it has a special metabolic pattern and immune microenvironment as represent by its lipophilic, transparent and metabolic pattern (35). Although the TME of ccRCC is infiltrated with large numbers of immune cells, a typical feature is that patients with high-degree CD4^+^ and CD8^+^T cell infiltration have a poorer prognosis (36). Secondly, the heterogeneity of renal cancer is reflected by the fact that ccRCC patients with a similar stage and grade may present completely different therapeutic response and OS after receiving the same treatment. They may present distinctive genetic and molecular alterations, experience different clinical courses, and exhibit different responses to the same therapy (37). It is therefore urgent to develop a comprehensive and robust ccRCC molecular typing model to help make specific personalized treatment plans.

In the present study, we made a comprehensive exploration of multiple omics including mRNA profiles, lncRNA profiles, miRNA profiles, DNA methylation profiles and simple somatic mutation to construct a subtyping tool to help address this important clinical issue. Through clustering the multi-omics data of 258 ccRCC patients after filtration, we generated two subtypes of ccRCC and found significantly different landscapes of mRNA, lncRNA, miRNA, DNA methylation, somatic mutation and CNVs between two subtypes of ccRCC patients. It was found that ccRCC patients in CS2 subtype group showed better prognosis than those in CS1 subtype group. In addition, pseudo-timing analysis further validated the robustness of our subtyping method by two distinct clusters and each of them was further stratified into four subtypes by the risk-related mRNA landscape. With the maturation of sequencing technology and the improvement of tumor atlas plans in recent years, more molecular typing studies on renal cancer have emerged. The TCGA team (37) remodeled ccRCC through a stratification remodeling study of from the levels of genomic alterations, DNA methylation profiles, RNA and proteomic signature, and finally found that PI(3)K/AKT pathway was recurrently mutated, suggesting that this pathway is a potential therapeutic target. On this basis, numerous ccRCC typing studies have emerged by using a single omics profile or specific gene set. Although these classification studies have to some extent provided new directions for the classification and precise treatment of renal cancer to some extent, they have their respective shortcomings. For example, Chen et al (39) integrated multiple-omics data of all RCC cases only based on one algorithm (COCA) without incorporating the lncRNA profile into the typing study. Besides, the classification methods used in these studies were relatively simple. Riketts et al (40) only integrated 843 RCC multiple-omics data and then categorized the data from each omics data alone, which did not achieve the true sense of multiple sets of data. The methods used in these classification studies can hardly be applied to clinical practice because they lack accuracy. In contrast, the current study used a variety of mainstream clustering algorithms to integrate multiple profiles at the same time to perform a classification study on ccRCC patients by paying special attention to analysis at each omics level one by one and the heterogeneity of the two groups of patients in TME. Surprisingly, this renal cancer subtyping model constructed according to differential biomarkers showed good survival prediction in both training and validation sets.

Importantly, our study extended previous studies in patient subtyping based on simple clustering analysis. The discrete subtype information is unable to capture inter- and intra-cluster relationships and provide the overall structure of patient distribution. To remedy these shortcomings, we applied graph learning approaches to uncover the tree structures of expression profiles among patients, believing that they can provide complementary information for clustering analysis and offer new insight into the complex landscape of ccRCC. Based on the pseudo-chronological analysis, we found that there may be a certain evolutionary relationship between CS1 and CS2. As the prognosis of the two subtypes is completely different, we performed a pseudotime analysis and found that the CS2 subtype was located at the beginning of the differentiation axis, while the CS1 subtype was in evolution. At the end of the axis, combined with the results of the prognostic analysis in previous studies, we speculate that the evolution of profiles between CS1 and CS2 can be used to explain the progressive condition of ccRCC patients. Among them, the up- and down-regulated genes that participated in the progression from the CS2 state to the S1 state may be the hub genes and potential therapeutic targets for ccRCC patients.

Enrichment analysis showed that extracellular matrix organization, cornification, keratinocyte differentiation and keratinization were distinctly enriched in CS1 subgroup was distinctly enriched in CS1 subgroup, which is in line with the following immune cell infiltration analysis. In GSVA analysis, we found that hypoxia, coagulation and myogenesis pathways were upregulated in CS1 subtype, while G2M_checkpoint, fatty acid metabolism and TGF β signaling onco-pathways were unregulated in CS2 subtype. All the GSVA results are consistent with the findings in the previously reported studies. Hypoxia is a typical character of RCC, which plays a crucial role in the development and progression of RCC together with angiogenesis and glucose metabolism, like a loop that is self-feeding(38). In this study, we found significant differences in PBRM1 and BAP1 mutation between the two subgroups. PBRM1 is the second most common mutated gene in ccRCC after VHL, as well as a component of the SWI/SNF chromatin remodeling complex. Different studies investigated the biological consequences and the potential role of PBRM1 alteration in RCC prognosis and identified it as a drug response modulator, although some results are controversial (39). The BAP1 gene has emerged as a major tumor suppressor mutated with various frequencies in numerous human malignancies, including uveal melanoma, malignant pleural mesothelioma, ccRCC, intrahepatic cholangiocarcinoma, hepatocellular carcinoma, and thymic epithelial tumor (40). BAP1 orchestrates chromatin-associated processes including gene expression, DNA replication, and DNA repair (41). BAP1 also exerts cytoplasmic functions, notably in regulating Ca2+ signaling at the endoplasmic reticulum(42). Taken together, we speculate that CS1 subtype of ccRCC tends to be infiltrated with more immune cells with apparent fibrosis or more mesenchyma as compared with CS2 subtype, which seems to be a more ‘clear cell-like’ classic renal carcinoma.

In recent years, immunotherapy has become one of the mainstays of treatment for renal cancer, and been recommended by the American Food and Drug Administration (FDA) as the first-line treatment for advanced renal cancer(43, 44). However, there is still a lack of targeted markers for predicting the benefit of immunotherapy in RCC patients. The TME contains a variety of components such as vascular interstitial components, chemokines and inflammatory factors (45). The composition and proportion of these components integrally determine the effectiveness of immunotherapy and directly determine the efficacy of anti-tumor immunity in renal cancer (46). In the present study, it was obvious that the proportion of different types of T cells in CS1 subtype was higher than that in CS2 subtype. They included activated CD4+T cells, activated CD8+T cells, regulatory T cells, T follicular helper cells and type 1 T helper cells. Subsequent GSVA analysis also demonstrated that immune cell functions were mainly enriched in CS1 subtype of ccRCC. However, although the infiltration scores of various immune cells in CS1 group were relatively higher, the prognosis of patients in CS1 group was poorer than that in CS2 group. In addition, the TIDE algorithm revealed that the immunotherapy benefit rate in CS2 group was significantly higher than that in CS1 group. Besides, the expression of immune checkpoints was significantly upregulated in CS1 subtype. All these results indicate that immune cell infiltration in CS1 subtype of ccRCC may play a co-inhibitory role, while the immune microenvironment in CS2 subtype demonstrated a better anti-tumor effect. This hypothesis is in line with previous studies. ccRCC is generally recognized as a immunogenic tumor, but it is also known for its immune escaping through regulatory T cells by upregulating a variety of inhibitory functions (47). Although patients with advanced renal cancer have more immune cell infiltration, these tumors often have immune escape. It is also proved that the degree of CD8+T cell infiltration in renal cancer is inversely proportional to the patient’s prognosis, meaning that a higher degree of CD8+T cell infiltration predicts a worse prognosis of patients with renal cancer (48).

It was found in our study that ccRCC patients of the two subtypes exhibited a significantly different sensitivity to molecular targeted drugs: patients in CS1 high-risk subgroup showed a higher sensitivity to Imatinib and Crizotinib as compared with those in CS2 subgroup, while patients in CS2 subgroup showed a higher sensitivity to Sracatinib, Listinib, Gefitinib and Dasatinib. To be noted, the response rate to immune checkpoint agents in ccRCC patients of CS2 subtype was higher than that in those of CS1 subtype. However, these immunotherapy markers are commonly used in other tumor types (such as PD1 and LAG3), and TMB was not identified as sensitive potential therapeutical target in ccRCC patients. Over the past 20 years, medical treatments for RCC have transitioned from non-specific immune approaches to targeted therapy against vascular endothelia growth factor (VEGF) and today’s novel immunotherapies. There are multiple targets, including platelet-derived growth factor and related receptors, and other inhibitors including mammalian target of rapamycin and the EMT and AXL tyrosine-protein kinase receptors. Nowadays, several immune-checkpoint inhibitors have been approved based on their significant curative effects in advanced RCC patients.

However, most other studies have focused on exploiting biomarkers for predicting the sensitivity of patients to immunotherapy and the optimal combination or selection of the existing agents. In the present study, we surprisingly noticed that CS1 subtype was insensitive to most molecular targeted drugs and immune inhibitory agents, which urged us to explore new potential drugs that could more effectively treat CS1 subtype. Also amazingly, we discovered a variety of potential drugs specific to this high-risk subtype. Among them, compound CGP.082996 was found to be associated with the MYC locus, suggesting that it is a potent antiproliferative agent against retinoblastoma (Rb)-positive tumor cells by exclusively arresting cells at G1 and reducing phospho-Ser 780 /Ser 795 on the Rb protein (49). CMK could inhibit the proliferation of lung adenocarcinoma cells both in vivo and vitro by targeting the RSK family members (50). JNJ-26854165 could induce wild-type p53- and E2F1-mediated apoptosis in acute myeloid and lymphoid leukemias (51). ZM 447439 is a novel promising aurora kinase inhibitor which can provoke antiproliferative and pro-apoptotic effects against gastroenteropancreatic neuroendocrine tumor diseases either alone or in combination with bio- and chemotherapeutic agents (52). GNF-2 can affect structural dynamics of the ATP-binding site by binding to the myristate-binding site of Abl (53). LFM-A13 exhibited a favorable pharmacokinetic behavior that was not adversely affected by the standard chemotherapy drugs vincristine, methylprednisolone, or l-asparaginase (when used as combination treatment, VPL) and significantly improved the chemotherapy response and survival outcome of mice challenged with BCL-1 leukemia cells (54). RO.3306 could enhance p53-mediated Bax activation and mitochondrial apoptosis in AML by inhibiting cyclin-dependent kinase 1(55). AZD-6482 is a specific PI3Kβ inhibitor that exerts an antitumor effect by inhibiting proliferation and inducing apoptosis of human glioma cells(56). Some of the potential drugs explored in the current study have not yet been reported before, such as AZD-6482, which is an intravenous platelet aggregation inhibitor which was reported to be used for the treatment of thrombosis but not cancer. It is worth-noting that CGP-60474, an inhibitor of cyclin-dependent kinase, has proved to be the most potential drug in that it alleviated tumor necrosis factor-α (TNF-α) and interleukin-6 (IL-6) in activated macrophages by downregulating the NF-κB activity and reducing the mortality rate in LPS-induced endotoxemia mice (57). In addition, VX.680, a highly potent and selective small-molecule inhibitor of Aurora kinases, could block cell-cycle progression and induce apoptosis in a diverse range of human tumor types including leukemia, colon and pancreatic tumors (58). Given that CS1 was more sensitive to the above-mentioned drugs and some drugs have achieved certain effects in clinical trials of some solid tumors, the above-mentioned drugs may provide new potential therapeutic targets for the high-risk subgroup of RCC.

This study provides new insights into molecular subtyping of ccRCC, and at the same time constructs a molecular prognostic model based on mRNA expression. However, this study has some limitations. Firstly, all the classification data used in this study include multi-omics data. In real clinical work, obtaining so many omics data from patients is expensive and time-consuming, so it is not easy for use in clinical practice. Secondly, although we compared drug sensitivities between different subgroups, specific experiments are still in need for further validation. Finally, the data used in this study are all from Western countries, and whether this model is applicable to other races and ethnicities remains to be determined. As the current clinical multi-omics panels are not yet popular, a risk model based on mRNA expression and the characteristics of subgroups needs to offer implications for the clinical work.

In summary, this is an innovative study on ccRCC molecular typing based on multiple omics, which describes differences between subgroups at multiple omics levels. As this prognostic model could efficiently predict the long-term survival and drug selection of ccRCC patients, it might bring new hope to the diagnosis and treatment of ccRCC.

## Data Availability

The data used to support the findings of this study have been deposited in the TCGA repository (https://portal.gdc.cancer.gov/) and ICGC repository (https://dcc.icgc.org/), respectively.

## Author contributions

Aimin Jiang, Yewei Bao and Anbang Wang have contributed equally to this work. Jie Wang, Bing Liu, Juan Lu and Linhui Wang conceptualized and designed this study. Xinxin Gan, Wenliang Gong, Zhenjie Wu and Yi Bao wrote the first draft of the manuscript. All authors contributed to the article and approved the submitted version.

## Acknowledgments

The authors sincerely thank all participants involved in this study. This study was funded by the National Natural Science Foundation of China (82072812, 81730073, 81872074), Shanghai Sailing Program (19YF1448300) and Clinical science and technology innovation project of Shanghai Shenkang Hospital Development Center (SHDC12018108).

## Supplementary Figures

Figure S1

The workflow of this study.

Figure S2

(A) Identification of the optimal cluster number by calculating CPI (blue line) and Gaps-statistics (red line) in TCGA-KIRC cohort.

(B) Consensus heatmap based on results from 10 multi-omics integrative clustering algorithms with the cluster number of 2, 3, 4 and 5, and related quantification of sample similarity using the Silhoutte score based on the consensus ensembles result.

Figure S3

Heatmap of subtype-specific upregulated and downregulated biomarkers using DEseq2 for the two identified subtypes in TCGA-KIRC cohort.

Figure S4

Heatmap of the immune related gene family of chemokines and receptors, co-inhibitors, costimulators, interferons and receptors and MHC expression in CS1 and CS2 subgroups.

Figure S5

(A) DNA substitution types including transition (Ti) and transversion (Tv) among CS1 and CS2.

(B) The most significant difference of mutated genes between CS1 and CS2.

(C) The lollipop plot illustrates the differential distribution of variants for PBRM1.

(D) Kaplan-Meier curves show the independent relevance between overall survival and PBRM1 mutation in CS1 and CS2 subgroups.

(E) The lollipop plot illustrates the differential distribution of variants for PBRM1.

Figure S6

The structure tomographs of the ten candidate small-molecule drugs for CS1 subgroup.

Figure S7

(A) Use of Boruta algorithm to calculate variable importance score (green is important variable, blue is shadow variable)

(B) The importance of the variables is sorted: from right to left, the importance is decreasing in order

## Supplementary Tables

Supplement Table 1 Biomarker signatures of each subtype.

Supplement Table 2 Recurrently amplified and deleted regions in the two subgroups calculated by GISTIC2.0

Supplement Table3 Name of discovered 53 kinds of small-molecule drugs that could be used as potential drugs for the treatment of CS1.

