## Supplementary figures and images for "Establishment of a Prognostic Prediction and Drug Selection Model for Patients with Clear Cell Renal Cell Carcinoma by Multi-Omics Data Analysis"

### Supplementary Figures S1_S7

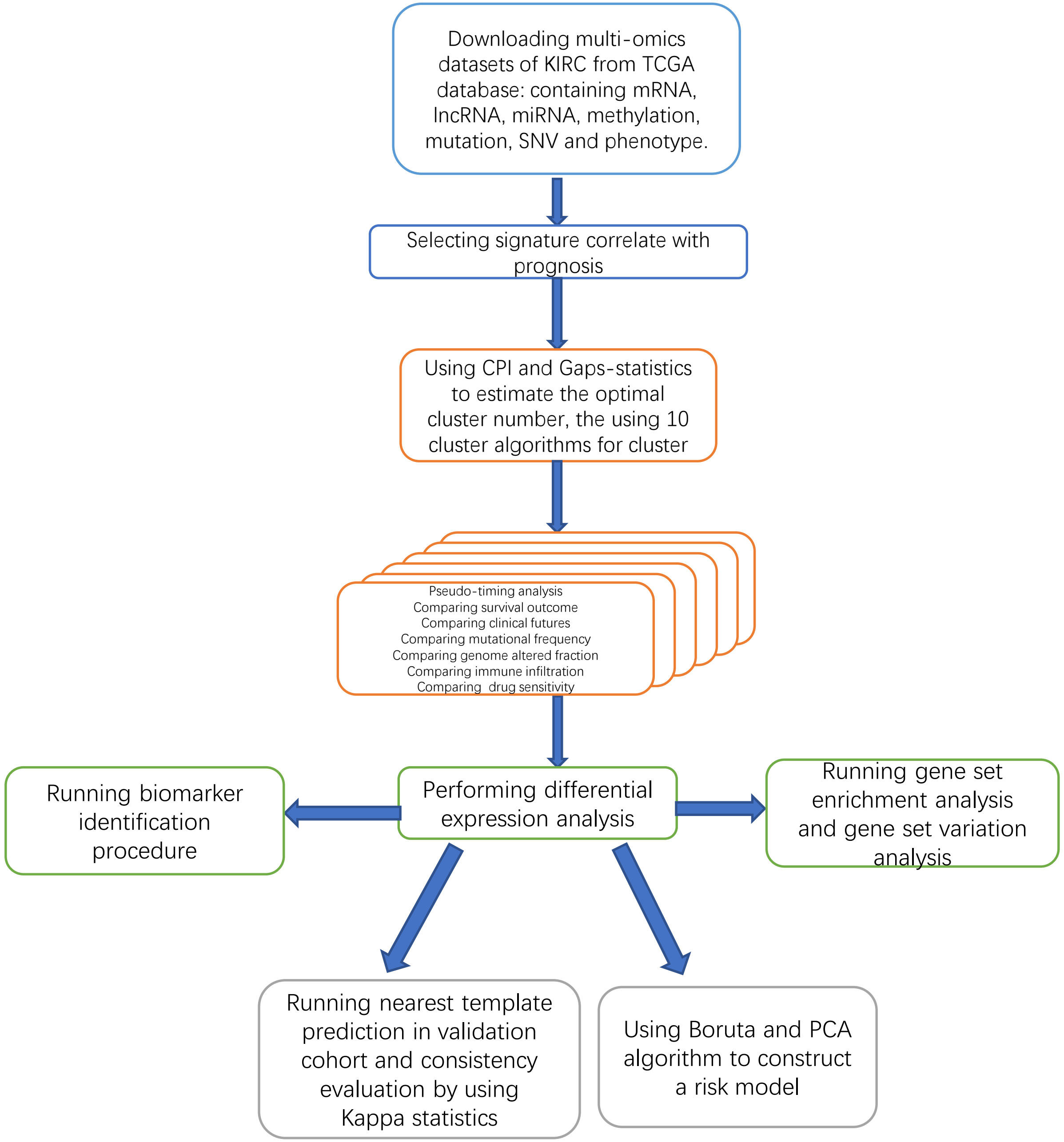

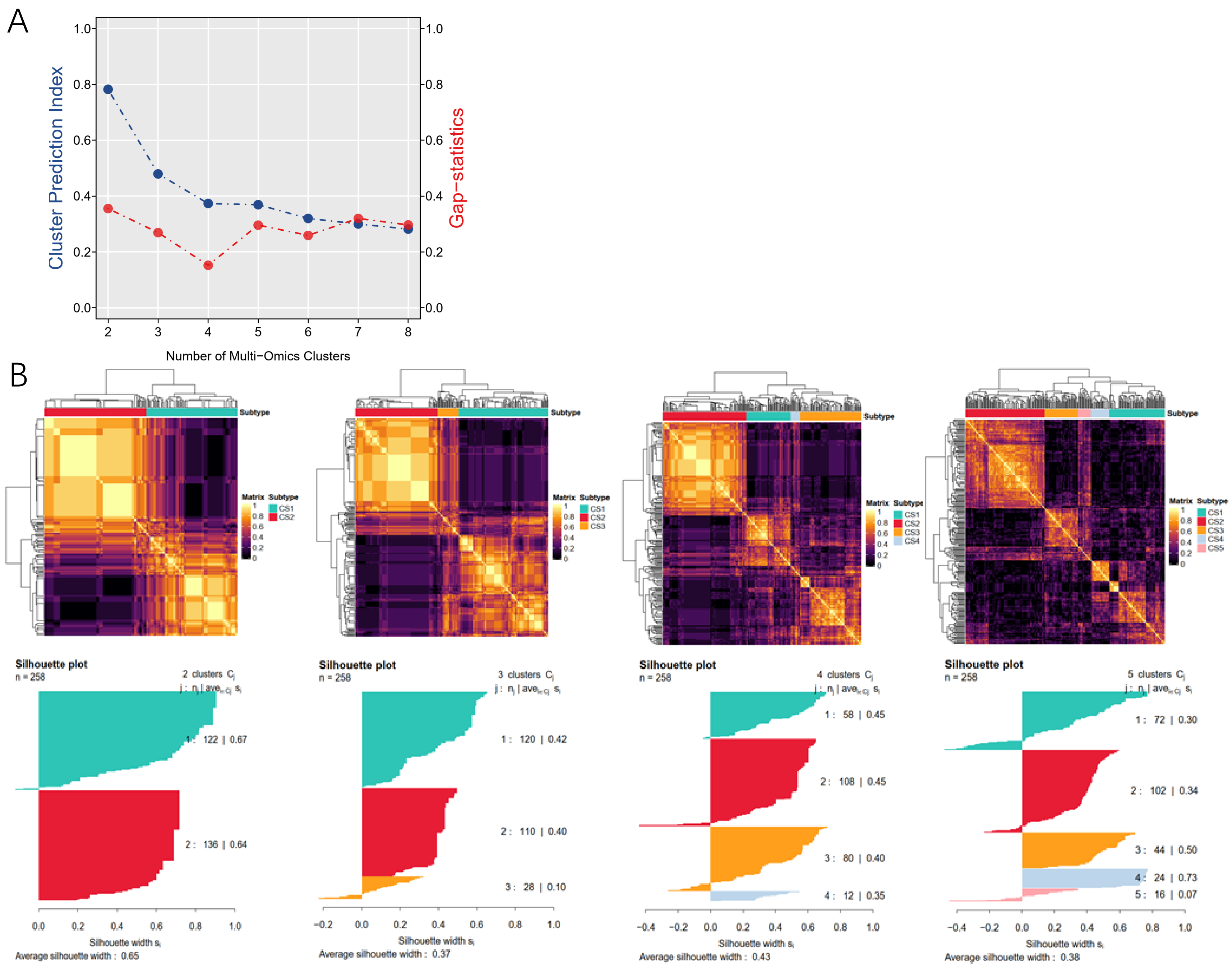

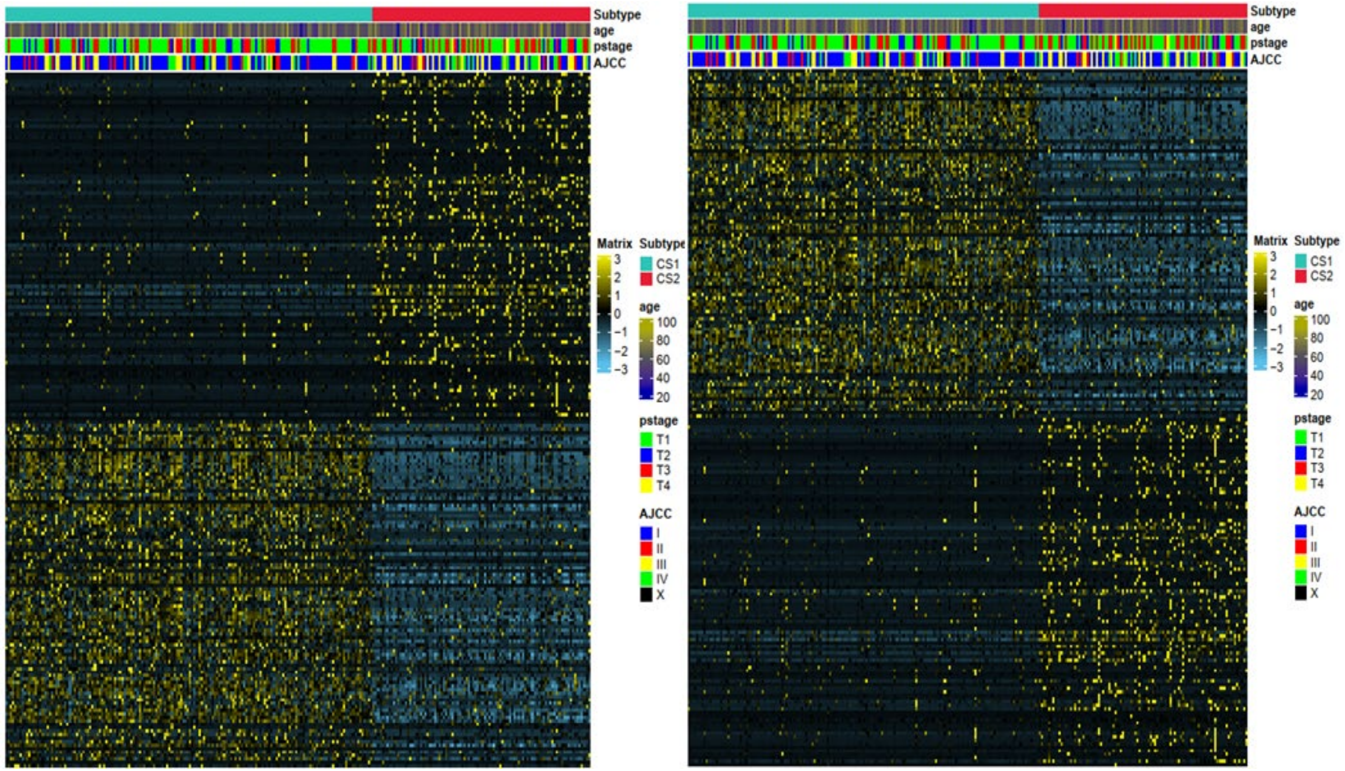

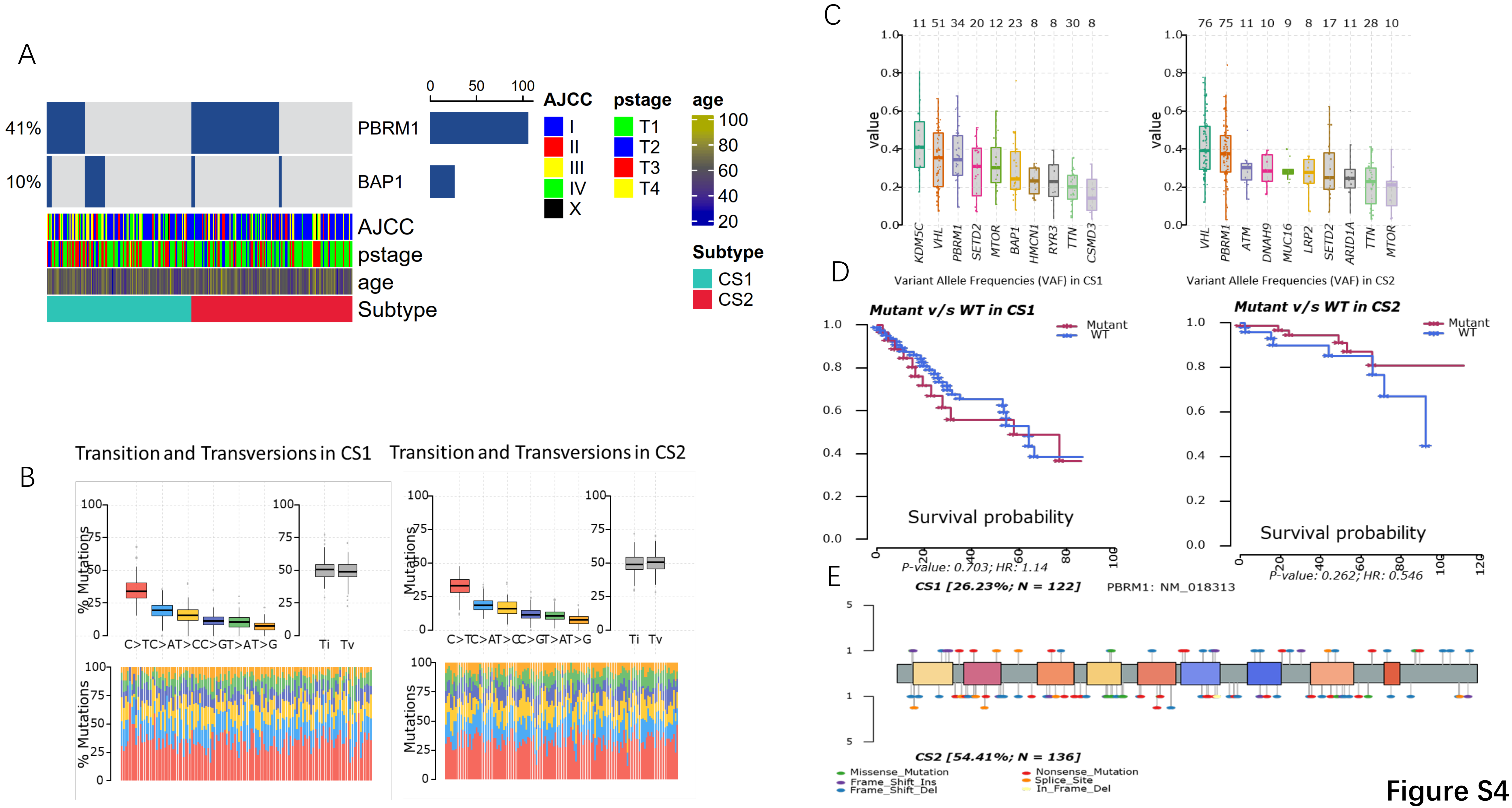

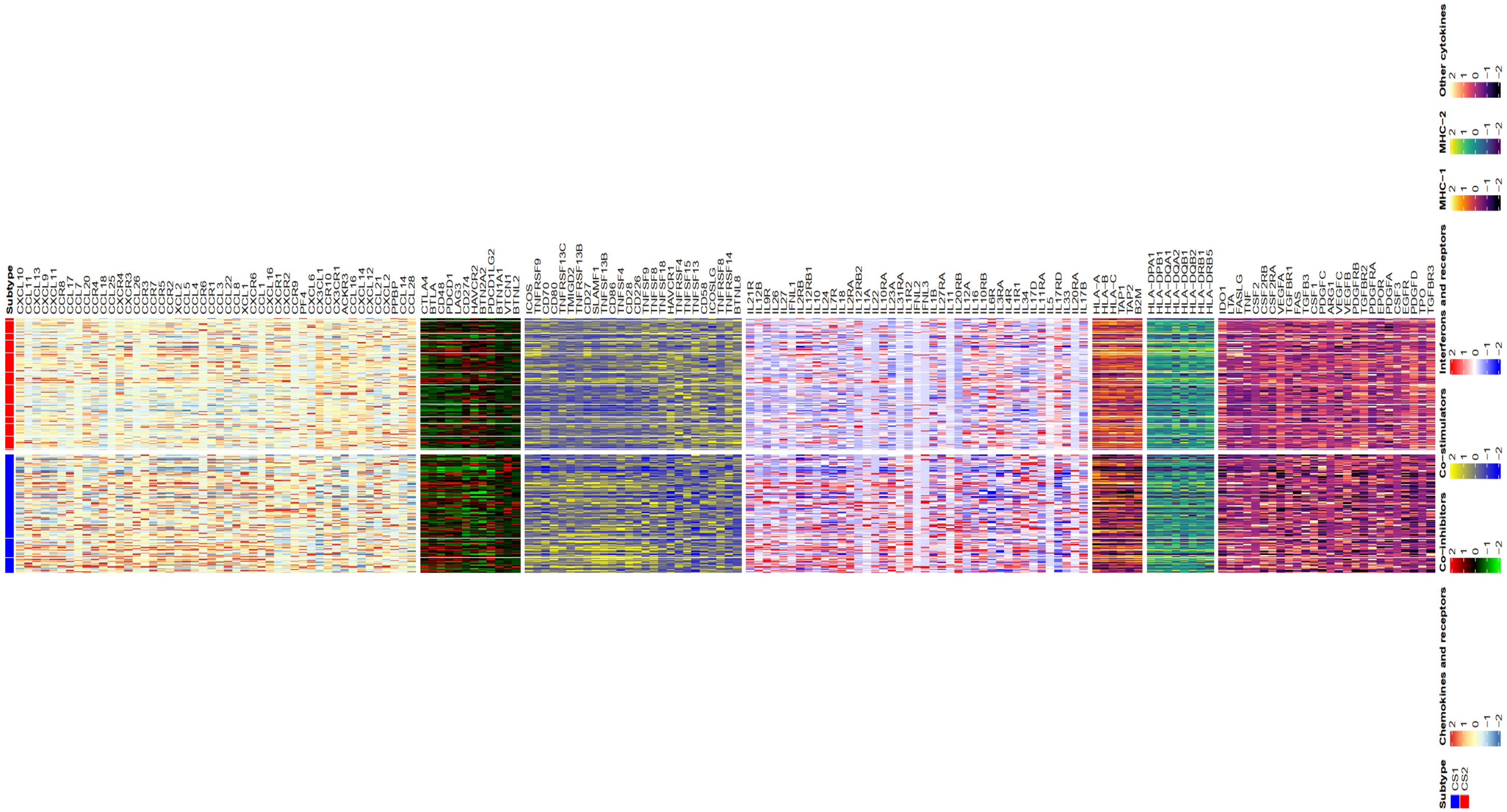

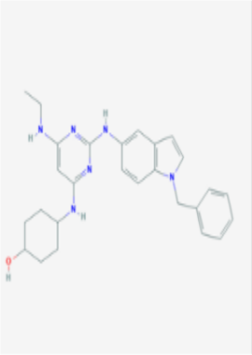

CGP.082996

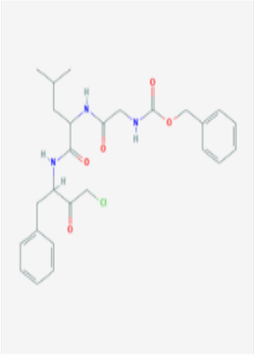

CMK

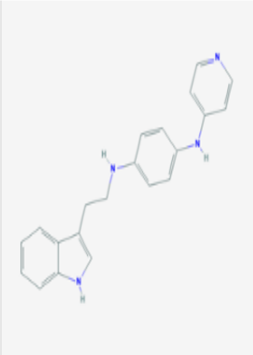

JNJ.26854165

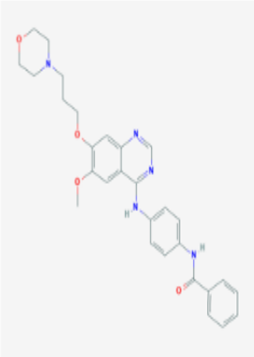

ZM.447439

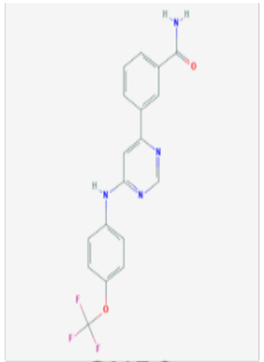

GNF.2

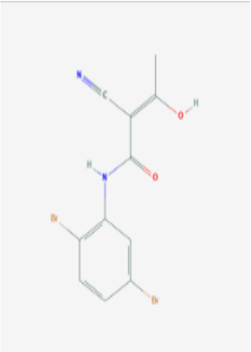

LAF.A13

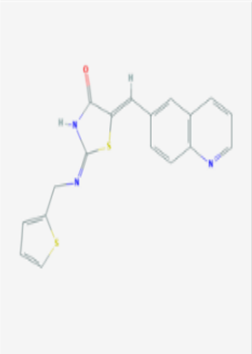

RO.3306

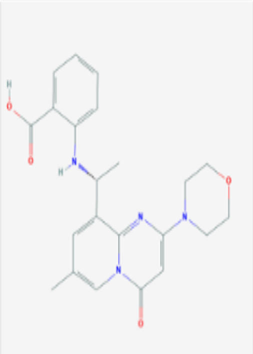

WO2009093972

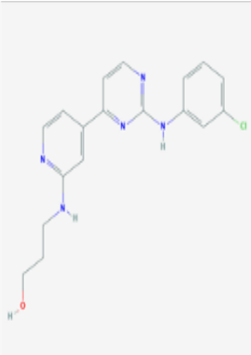

CGP.60474

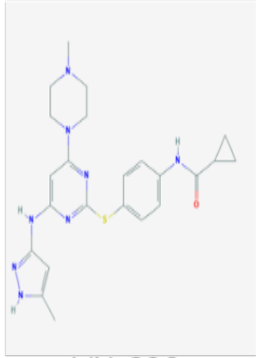

VX.680

**Figure S6**

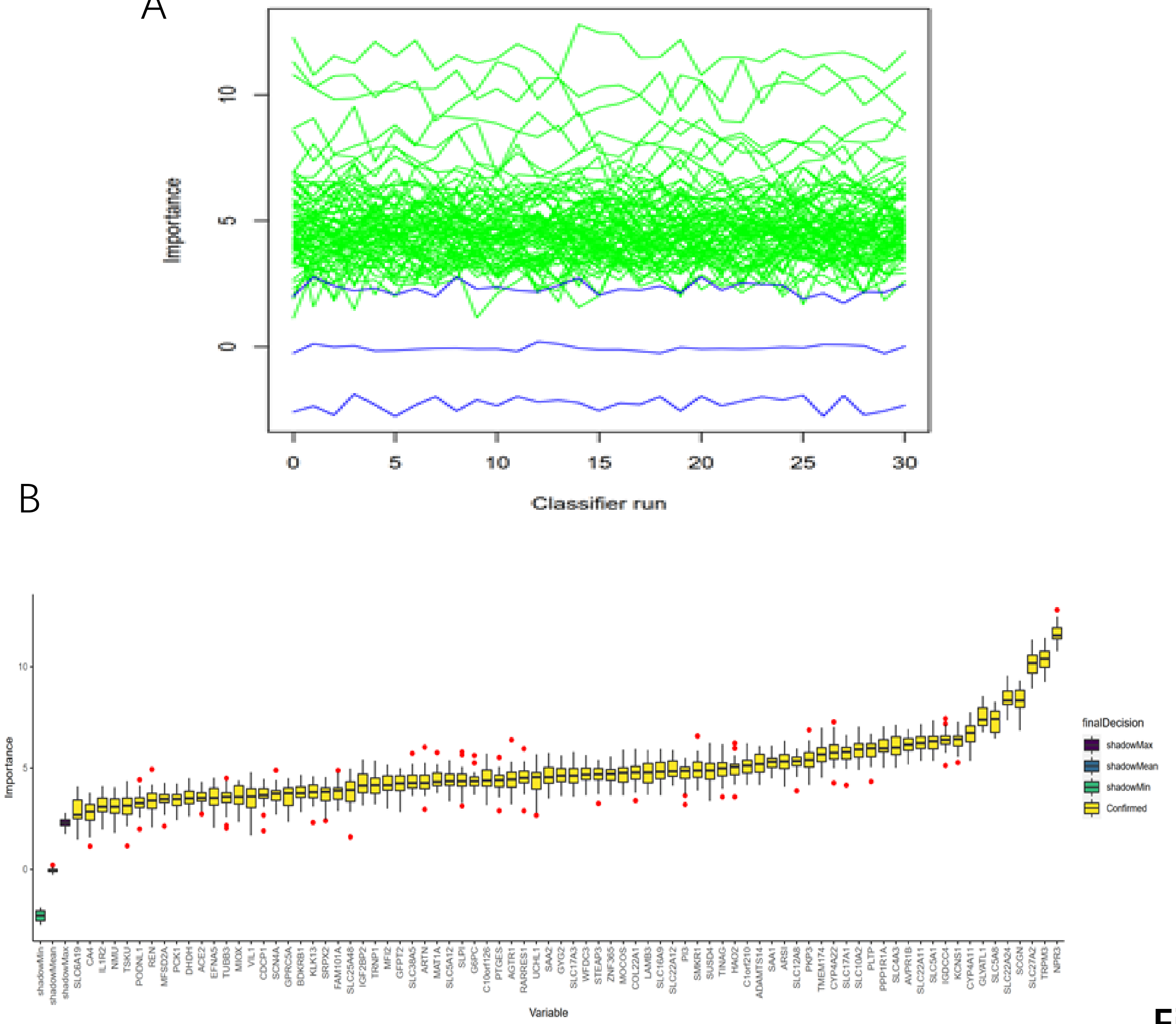
