## Supplementary material for "Establishment of a Prognostic Prediction and Drug Selection Model for Patients with Clear Cell Renal Cell Carcinoma by Multi-Omics Data Analysis": table 1

*Summary descriptives table by groups of `cluster'*

| | CS1<br>*N=122* | CS2<br>*N=136* | p.overall |
|---|---|---|---|
| fustat: | | | <0.001 |
| 0 | 80 (65.6%) | 117 (86.0%) | |
| 1 | 42 (34.4%) | 19 (14.0%) | |
| futime | 1212 (984) | 1506 (1131) | 0.026 |
| AJCC: | | | 0.083 |
| I | 58 (47.5%) | 86 (63.2%) | |
| II | 12 (9.84%) | 13 (9.56%) | |
| III | 29 (23.8%) | 22 (16.2%) | |
| IV | 22 (18.0%) | 14 (10.3%) | |
| X | 1 (0.82%) | 1 (0.74%) | |
| age | 60.7 (11.7) | 60.8 (12.2) | 0.970 |
| PFI: | | | 0.005 |
| 0 | 79 (64.8%) | 110 (80.9%) | |
| 1 | 43 (35.2%) | 26 (19.1%) | |
| PFI.time | 1038 (970) | 1290 (1041) | 0.045 |
| gender: | | | 0.214 |
| FEMALE | 41 (33.6%) | 57 (41.9%) | |
| MALE | 81 (66.4%) | 79 (58.1%) | |
| grade: | | | <0.001 |
| G1 | 2 (1.64%) | 7 (5.22%) | |
| G2 | 50 (41.0%) | 68 (50.7%) | |
| G3 | 41 (33.6%) | 51 (38.1%) | |
| G4 | 28 (23.0%) | 8 (5.97%) | |
| GX | 1 (0.82%) | 0 (0.00%) | |
| laterality: | | | 0.658 |
| Bilateral | 1 (0.82%) | 0 (0.00%) | |
| Left | 55 (45.1%) | 59 (43.4%) | |
| Right | 66 (54.1%) | 77 (56.6%) | |
